# The Brain Encyclopedia Atlas Project (BEAP): A Literature-Synthesis-Derived Functional Atlas of the Human Brain

**DOI:** 10.64898/2026.05.05.722843

**Authors:** Oren Poliva

## Abstract

The neuroscience literature contains thousands of studies linking cognitive, sensory, and motor functions to specific brain regions, yet this knowledge remains fragmented across experimental modalities, naming conventions, and spatial reference systems. Consequently, relating reported activations, lesions, or stimulation sites to prior findings often requires substantial manual synthesis. The Brain Encyclopedia Atlas Project (BEAP) was developed to address this challenge by providing a spatially grounded neuroinformatics framework for organizing literature-defined functional neuroanatomy.

BEAP is an expert-curated resource that aggregates and spatially indexes literature-defined cortical and subcortical regions within a common anatomical reference framework. The project identifies 106 neocortical fields and 18 cerebellar fields through analysis of published figures from 1,465 human studies using functional neuroimaging, intracranial electrophysiology, cortical stimulation and lesion mapping studies. These regions were manually aligned to standard anatomical templates and associated with parcels of the Human Connectome Project multimodal parcellation (MMP1). Inclusion criteria required convergent functional evidence, lesion support, and boundary-related contrasts. In addition, putative connectivity profiles were compiled from 475 non-human primate tract-tracing studies through comparison with established macaque atlases and homology-based mapping to human cortical territories. Beyond the neocortex, 340 allocortical, diencephalic, cerebellar, and brainstem nuclei were delineated through comparison with histological atlases and related anatomical studies.

The resource is publicly accessible through an interactive three-dimensional brain atlas linked directly to a curated encyclopedia of functional neuroanatomy. Each entry synthesizes functional descriptions, boundary-defining evidence, lesion associations, internal organization, alternative nomenclature, and putative connectivity annotations grounded in the primary literature. BEAP further supports research interoperability through downloadable volumetric and surface-based atlas files compatible with standard neuroimaging environments, including FSLeyes and FreeView.

By integrating heterogeneous literature-defined functional territories into a common spatial reference framework, BEAP provides a practical resource for contextualizing neuroimaging findings, supporting literature synthesis, and facilitating exploration of human functional neuroanatomy.

## INTRODUCTION

Since the advent of functional magnetic resonance imaging (fMRI) more than three decades ago, the neuroscience literature has accumulated thousands of reports describing functionally specialized regions of the human brain. These include widely recognized territories such as the primary and extrastriate visual cortices, the frontal eye field, Broca’s area, and the fusiform face area. The literature also describes less commonly referenced regions, including the inferior frontal face area, the dorsal precentral speech area, and Exner’s area. Many of these regions recur across task-based imaging, lesion studies, stimulation experiments, and intracranial recordings. Collectively, this body of work has produced a rich but highly fragmented body of functional localization evidence distributed across experimental modalities, historical traditions, and naming conventions.

Existing human brain atlases only partially address this problem. Many commonly used atlases are organized around anatomical landmarks, cytoarchitecture, receptor architecture, or molecular properties rather than the literature-defined functional entities emphasized in experimental neuroscience [1]. Coordinate-based meta-analytic platforms such as Neurosynth and NeuroQuery provide large-scale statistical synthesis of functional neuroimaging studies but are largely limited to coordinate-reported fMRI datasets and do not preserve the spatial context present in published figures, including activation extent, relationships to sulcal landmarks, and boundary contrasts between neighboring regions. In addition, these resources generally do not integrate lesion studies, intracranial electrophysiology, cortical stimulation findings, or tract-tracing evidence from the non-human primate literature.

A major exception among contemporary atlas frameworks is the Human Connectome Project multimodal parcellation atlas (MMP1) [2], which delineates 180 cortical parcels per hemisphere using convergent MRI-derived measurements, including cortical thickness, relative myelin density, resting-state connectivity, and task-fMRI data. Due to its widespread adoption in neuroimaging research, MMP1 provides a practical reference framework for organizing functional neuroanatomical findings. Nevertheless, most MMP1 parcels are not accompanied by extensive literature-based functional syntheses, and relating parcel labels to the broader experimental literature often still requires substantial manual interpretation.

The present work introduces the Brain Encyclopedia Atlas Project (BEAP), an expert-curated neuroinformatics resource designed to organize literature-defined functional neuroanatomy within a common spatial reference framework using MMP1 as an indexing system. Rather than proposing a new anatomical parcellation, BEAP functions as a literature-synthesis and spatial-indexing framework that associates previously reported functional territories with one or more MMP1 parcels through figure-based localization and predefined curation rules. Spatial localization is therefore treated as an approximate parcel-level indexing procedure intended to relate heterogeneous findings to a shared anatomical framework rather than to define precise anatomical borders.

To construct the resource, published figures from functional neuroimaging, lesion mapping, intracranial electrophysiology, cortical stimulation, and related methodologies were manually aligned to standard anatomical templates and associated with MMP1 parcels. Human functional evidence was further supplemented with non-human primate tract-tracing studies to derive putative connectivity annotations, and with histology-based atlases to delineate subcortical and brainstem nuclei. The resulting platform integrates these data into an interactive three-dimensional atlas linked directly to encyclopedia-style entries summarizing functional descriptions, lesion associations, internal organization, boundary-defining evidence, alternative nomenclature, and putative connectivity relationships.

By providing a spatially grounded framework for organizing heterogeneous functional neuroanatomical findings, BEAP aims to facilitate interpretation of neuroimaging results, support literature synthesis, and provide a practical educational and research resource for human functional neuroanatomy.

## METHODS

### Overview

The Brain Encyclopedia Atlas Project (BEAP) is a whole-brain neuroinformatics resource designed to organize heterogeneous functional neuroanatomical evidence within a common spatial reference framework. The resource was constructed by translating figure-based localization reports from the neuroscience literature into standardized brain anatomical templates. Because figure-based localization requires expert interpretation, all curation decisions were guided by predefined decision rules intended to promote internal consistency and transparency.

Published figure panels and stereotaxic coordinates, when available, were aligned to canonical volumetric and surface templates and associated with parcels of the Human Connectome Project multimodal parcellation atlas (MMP1) [2]. Human functional and lesion evidence was supplemented with non-human primate tract-tracing studies to derive putative connectivity annotations, and with histology-based atlases to delineate subcortical and brainstem nuclei. The resulting evidence was aggregated into encyclopedia-style entries linked to an interactive three-dimensional atlas interface.

Because the primary objective of BEAP is to organize literature-defined functional knowledge, spatial localization was treated as an approximate indexing procedure rather than a precise anatomical measurement. Localization was therefore performed primarily at the parcel level and intended to capture the general cortical territory associated with a finding rather than its exact spatial extent.

### Literature identification and eligibility

A structured literature search strategy was conducted using Google Scholar to identify studies reporting associations between behavior and localized brain activity using functional brain mapping techniques, including functional magnetic resonance imaging (fMRI), positron emission tomography (PET), transcranial magnetic stimulation (TMS), magnetoencephalography (MEG), stereotactic electroencephalography (sEEG), electrocorticography (ECoG), direct electrical cortical stimulation, and lesion mapping.

Studies were eligible for inclusion only if they contained figure panels depicting the spatial location of the reported activation, lesion, stimulation site, or recording location. Coordinate-only studies were excluded because stereotaxic coordinates alone often lack information regarding spatial extent, anatomical context, and boundary relationships across studies with heterogeneous normalization and smoothing procedures.

Initial searches used established cortical field names as keywords. Additional studies were identified through citation tracing and relevant review articles. The search strategy was iterative, with alternative nomenclature encountered during screening subsequently incorporated into additional searches.

### Figure selection and coordinate handling

For each eligible study, the figure panel that most clearly depicted the reported spatial effect was selected for localization. When stereotaxic coordinates were reported, they were extracted to supplement figure-based localization but were not used as substitutes for figures. When figures and coordinates referred to the same cortical territory, both sources informed localization. In cases of discrepancy, figure-based localization was prioritized.

Talairach coordinates were converted to MNI space using the BioImage Suite online conversion tool (https://bioimagesuiteweb.github.io/webapp/mni2tal.html).

### Template matching and parcel identification

Published figures were compared against standard anatomical templates using widely adopted neuroimaging visualization software. Volumetric figures depicting coronal, axial, or sagittal slices were aligned to the MNI152 T1-weighted template using FSLeyes. Surface-based figures were compared to the fsaverage surface template using FreeView. Three-dimensional rendering figures were compared to a three-dimensional rendering of the MNI152 T1-weighted template using FreeView.

Alignment was guided by gross anatomical landmarks, including gyral and sulcal patterns, ventricular geometry, cortical poles, the central sulcus, and the sylvian fissure. Spatial correspondence was treated as approximate and resolved primarily at the level of atlas parcels.

MMP1 parcels were visualized using the original surface-based annotation files [2] (Fig.1A) or using a volumetric representation of the atlas generated using custom conversion scripts (Fig.1B). When anatomical correspondence was visually straightforward, parcel identification was performed directly within the visualization software. More ambiguous cases underwent additional manual alignment procedures.

**Figure 1.**
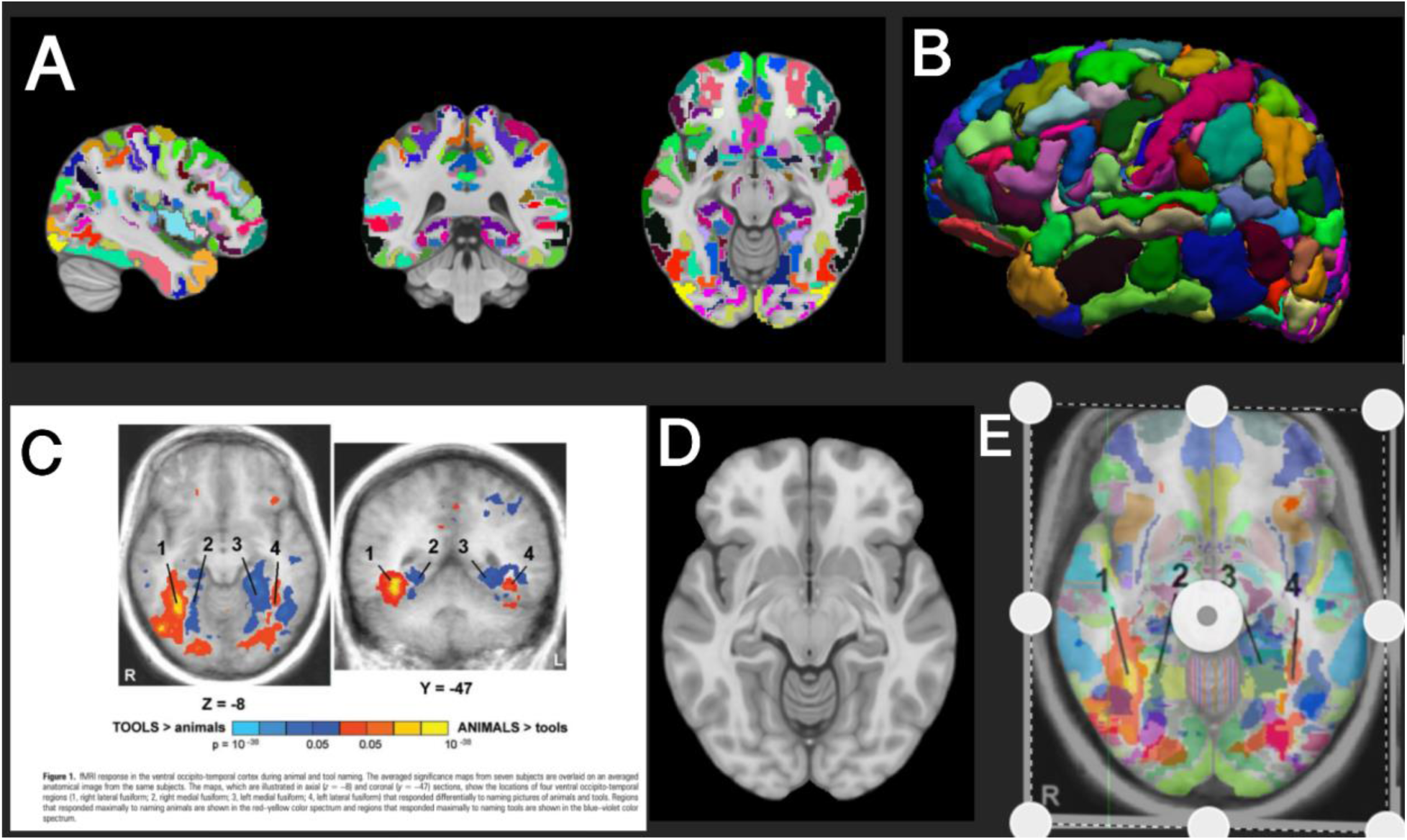
Workflow for associating MMP1 parcels with previously reported findings. Association of MMP1 parcels with previously published findings occurred in four stages. First, a custom script was used to convert the MMP1 atlas into volumetric space to enable comparison with slice-based figures from the literature (A,B). Second, figures from research articles reporting functional brain mapping results (fMRI, PET, ECoG, sEEG, MEG, TMS and cortical stimulation) were identified. As an example (C), we show a figure from Chao et al., [35] who reported significant BOLD activation on axial and coronal slices when participants viewed animals in contrast to tools. To estimate the location of this activation, a corresponding axial slice from the MNI152 template was selected based on anatomical similarity (D). The published figure was then superimposed on the template brain and manually aligned so that major anatomical landmarks matched (E). The MMP1 parcels underlying the activation were then identified and recorded. Figure adapted from Chao et al. (2002) with permission.

### Manual alignment procedure

To document correspondence between published figures and reference templates, a section of the template brain that most closely matched the published figure was first identified. For example, if a study presented an axial volumetric slice (e.g., Fig. 1C, left), a corresponding axial section from the MNI152 template was selected for comparison (Fig. 1D). The selected figure panels and the corresponding template image, with MMP1 parcels superimposed, were then imported into a graphics editing environment. Published figures were manually scaled and aligned to the reference template using major anatomical landmarks as anchors. The activation extent was subsequently traced on separate layers and associated with the underlying MMP1 parcels (Fig. 1E).

This procedure enabled each study to be associated with one or more parcels while preserving the approximate spatial extent represented in the original publication. As evidence accumulated across studies, overlapping parcel assignments were aggregated to represent candidate cortical fields.

### Lesion evidence integration

For lesion studies, figures depicting lesion extent were visually inspected and localized using the same template-matching procedure. Studies involving lesions spanning entire lobes or multiple lobes were excluded because of insufficient spatial specificity.

For eligible studies, lesion territories and associated behavioral deficits were documented and linked to overlapping BEAP regions. Lesion associations were recorded regardless of whether they supported or contradicted the putative function of the region, allowing entries to capture both convergent and conflicting evidence.

For voxel-based lesion-symptom mapping studies, statistically significant overlap clusters were used as localization targets.

### Cortical field designation criteria

A literature-defined cortical field was designated when the following criteria were satisfied (Fig. 2 for an example):

1. At least five functional studies associated the same parcel set with a common or closely related function.
2. At least one lesion study linked damage involving the same territory to disruption of the corresponding function.
3. For each border with a neighboring region, at least one functional study had to demonstrate a distinction between the candidate field and the adjacent territory, either through circumscribed activation confined to one region or through dissociable functional responses between the two regions.

**Figure 2.**
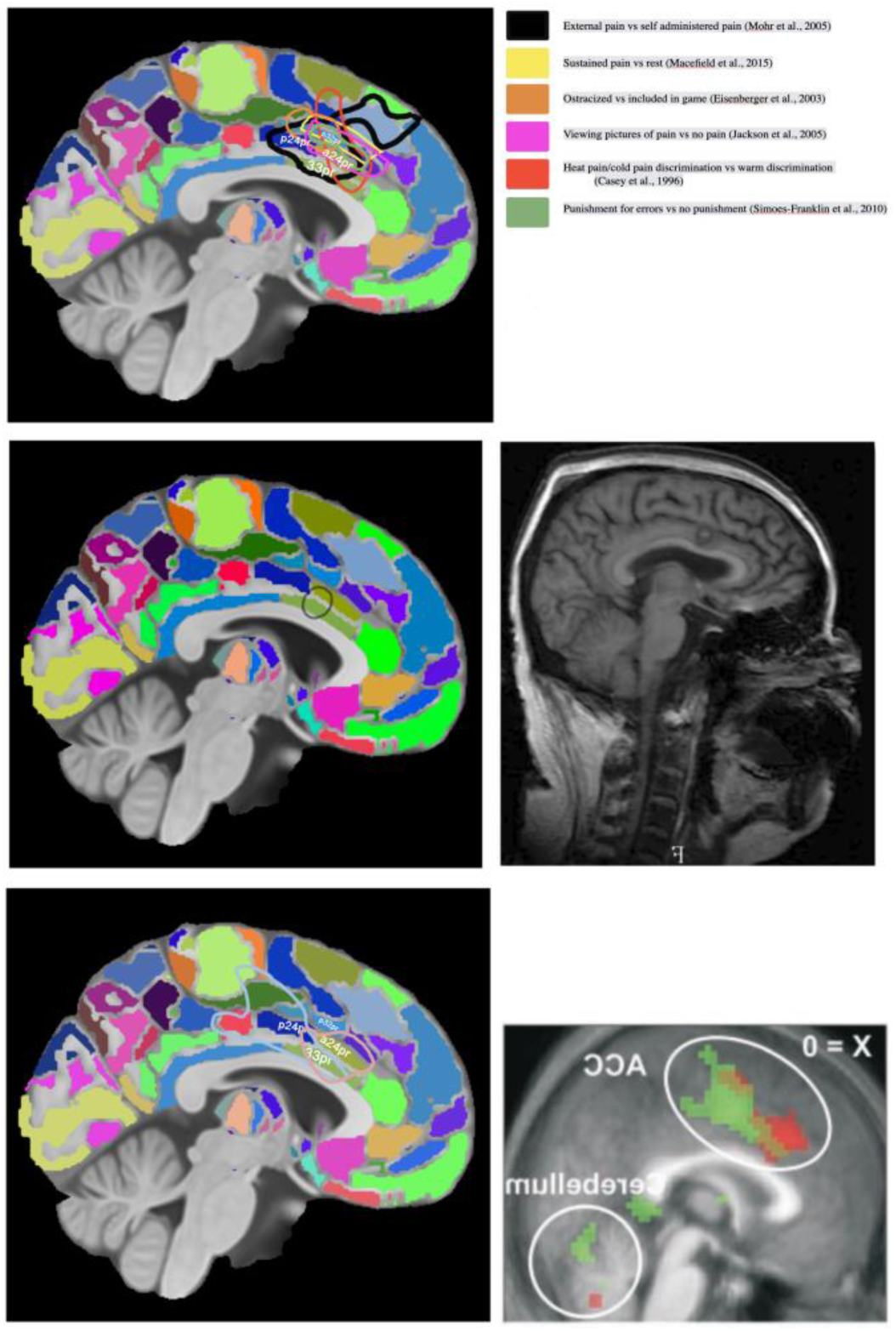
Criteria used to define a cortical field in the BEAP framework. Three criteria were used to identify an MMP1 parcel, or group of parcels, as a cortical field. First, at least five functional studies using brain mapping methods (fMRI, PET, ECoG, sEEG, TMS, MEG, or cortical stimulation) had to report activation within the same parcel set associated with the same or closely related function. An example is shown for anterior middle cingulate cortex (aMCC). Using the figure-matching pipeline, five fMRI studies and one PET study associated parcels a24pr and p32pr with the experience or anticipation of emotional and physical pain (top). Second, at least one lesion study had to link damage within the same territory to disruption or loss of the same function. As an example (middle), Yen et al., [36] reported chronic intractable pain in ten patients associated with tumors involving the middle cingulate cortex, with symptom improvement observed in six patients following tumor resection. Third, for each border of the region, there is at least one functional study, preferably fMRI because of its broader cortical coverage, which demonstrated a functional boundary with a neighboring region or circumscribed activation within one of the regions but not the other. In the example shown (bottom), Singer et al. [37] reported differential activation between parcel p24p, associated with first-hand pain, and parcel a24pr, associated with empathic responses when participants observed their partner receiving a mild electric shock.

These criteria were intended to prioritize conservative aggregation of evidence and reduce premature fragmentation of cortical territories. In practice, literature searches for a candidate cortical field were generally concluded once these predefined designation criteria had been satisfied. However, because many studies were relevant to multiple cortical territories, additional evidence continued to accumulate iteratively as neighboring regions and related functional systems were curated. Consequently, encyclopedia entries should not be interpreted as exhaustive systematic reviews, but rather as structured and continuously expandable syntheses of representative evidence.

### Expansion, merge, and subdivision rules

Expansion, merge, and subdivision decisions followed predefined rules intended to maintain consistency across the atlas.

1. Parcels that did not satisfy the minimal criteria to be designated as a cortical field but were strongly supported by non-human animal anatomical evidence, were retained as provisional entries and noted as requiring additional human evidence in the *Notes* section.
2. If a study extended a cortical field into an adjacent MMP1 parcel lacking an existing functional assignment, the field was expanded to include that parcel (i.e., there was no need for 5 studies to converge onto the same parcel).
3. If a study extended a cortical field into a neighboring cortical field with a distinct established function, the overlapping parcel was designated a transition region. This designation was documented in the *Internal Organization* section.
4. If fewer than five functional studies assigned a novel function to a cortical field with an already established different function, and the functional description could not reasonably be generalized to encompass both, the novel function and supporting evidence were documented in the *Notes* section.
5. If at least five functional studies reported a functionally distinct subregion within a field with consistent spatial localization, the field was split into two distinct cortical fields.
6. If at least five studies reported a functionally distinct subregion but with inconsistent spatial arrangements across studies, the region was considered to contain multiple distinct cortices supporting different functions.
7. If fewer than five studies reported functionally distinct subregions whose functions reflected nuanced variants of the broader field function, this internal organization was documented in the *Internal Organization* section.
8. If fewer than five studies reported evidence directly contradicting the established function of a region, the conflicting evidence was documented in the *Notes* section.
9. If a study assigned a novel function to a region and the functional description could be generalized to encompass both existing and novel roles, the description was updated to a more inclusive formulation.
10. If at least five functional studies and at least 1 lesion study supported a novel function incompatible with the established function, the region was treated as containing more than one cortical field.
11. Findings supporting a role for a cortex in one hemisphere were treated as evidence for this area also in the contralateral hemisphere, unless evidence indicated that the contralateral field is involved with a different function altogether.

### Putative connectivity annotations

Tract tracing is an anatomical method commonly used in non-human primates to delineate connectivity by identifying the regions that provide afferent input to a region of interest (retrograde tracing) and the regions that receive its efferent output (anterograde tracing) [3, 4]. This level of cellular and directional specificity is not directly captured by human neuroimaging measures such as functional connectivity or effective connectivity, which infer statistical coupling or directed interactions rather than tracing physical axonal pathways [5, 6]. Consequently, tract tracing is widely regarded as the gold-standard experimental approach for mapping anatomical connectivity.

In this study, the non-human primate tract-tracing literature was surveyed to compile putative connectivity profiles. A connection was annotated as present if it was reported in at least one tract-tracing study, even if other studies did not report that connection, reflecting differences in tracer type, injection site, and sampling across reports. Putative connectivity in humans was then hypothesized by assigning a macaque homologue to each human region based on established homologies (e.g., V1, AIP, FEF) or relative anatomical position with respect to regions with established homology. Each BEAP cortical field was compared to corresponding fields in two established macaque atlases, the M132 atlas (Fig. 3-top) and the Pandya atlas (Fig. 3-bottom).

**Figure 3.**
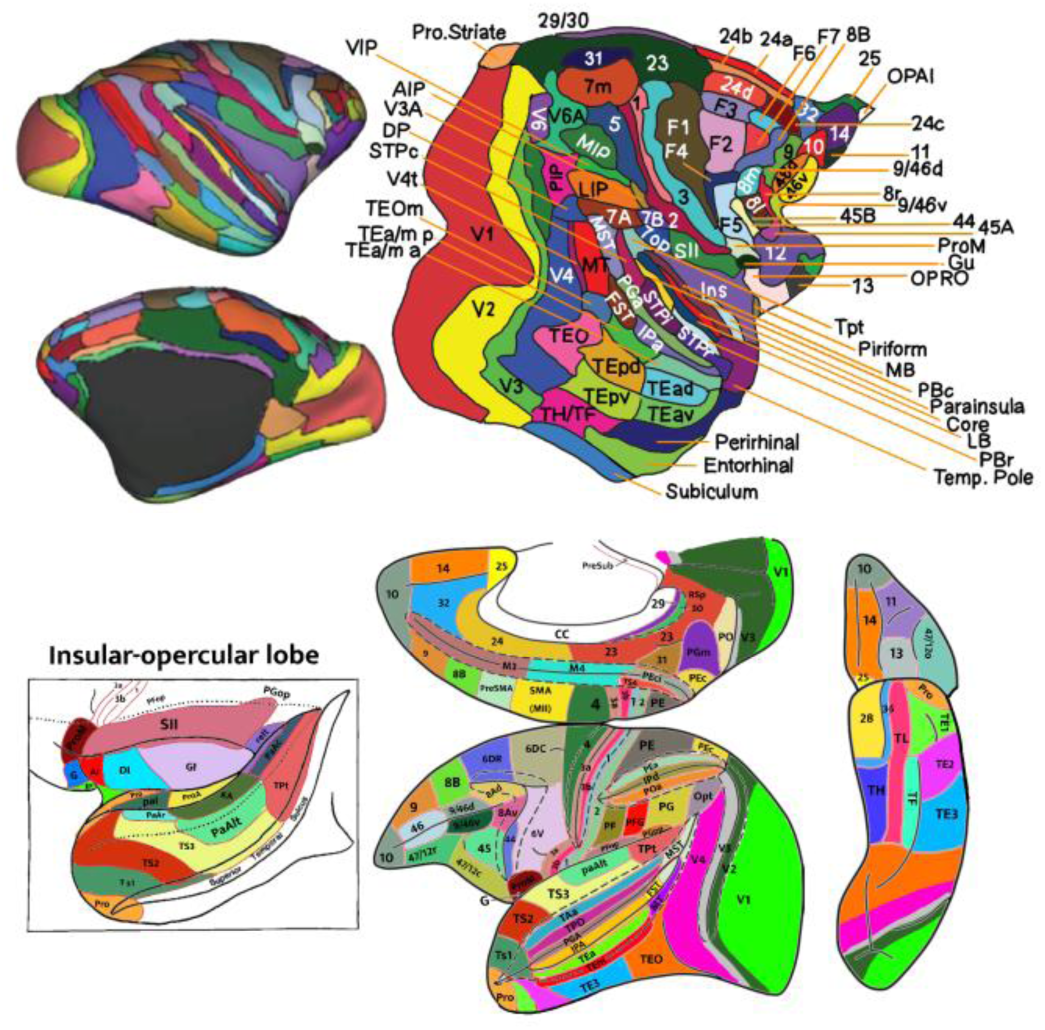
Assignment of putative connectivity through comparison with macaque atlases. Putative connectivity between human cortical fields was inferred by assigning candidate homologues in the macaque brain and examining their connectivity patterns in two macaque atlases commonly used in tract-tracing studies: the M132 atlas (top) [4] and the Pandya atlas (bottom) [12]. Figures were adapted with permission from the original sources.

When homology was uncertain, the rationale for the proposed mapping, including relevant references, was documented in the *Notes* section for that region. When available, human functional imaging or cortical electrophysiology studies reporting task-related co-activation or functional connectivity were also cited as supplementary context. These connectivity profiles are intended as qualitative reference annotations to support interpretation and hypothesis generation, not as direct evidence of human structural connectivity.

### Cerebellar field mapping

Cerebellar fields were mapped using procedures analogous to those applied to the cerebral cortex. Because cerebellar organization is relatively conserved across mammalian species [7], evidence from rodents and cats was also incorporated in addition to primate studies.

Functional assignments were based on figure localization, lesion evidence, connectivity studies, and functional neuroimaging reports. Resting-state functional connectivity studies linking cerebellar territories to cortical systems were used as organizational references for encyclopedia entries (see Fig.S1 in Supplemental Material 3).

### Subcortical and brainstem delineation

Unlike cortical fields, subcortical and brainstem nuclei were delineated primarily using histology-based atlases, reflecting differences in data availability and the greater anatomical stability of these structures. Primary delineation was based on cross-referencing two histology-based human brain atlases: *Atlas of the Human Brain* [8] and the *Allen Human Brain Atlas* [9]. When atlases agreed, nuclei were manually delineated directly on the MNI152 template using FSLeyes. When atlases disagreed or coverage was incomplete, additional peer-reviewed histological studies were consulted as tiebreakers (see Table I).

**Table I:** Literature used as tie-breakers for subcortical nuclei.

| Region | Reference |
| --- | --- |
| rhinencephalon | [17] |
| amygdala | [18] |
| entorhinal cortex | [19] |
| hippocampus | [20] |
| striatum | [21] |

For brainstem delineation, atlas plates from an established histological atlas [10-Chapter 10] were digitized and visually aligned to corresponding MNI152 template sections using ventricular and brainstem landmarks, and nuclei were manually traced in template space (Fig.S2-Top in Supplemental Material 3). Several structures not reliably visible on T1-weighted MRI were localized using the Big Brain Project, which is a histological atlas that is available directly in MNI152 space (Fig.S2-Bottom in Supplemental Material 3) [11]. Additional nuclei, which were not reported in any of the histological atlases, but are known from the literature, were delineated by integrating anatomical descriptions relative to known nuclei, from the mammalian literature (Table II). Due to the interpretive nature of this process, subcortical nuclei should be regarded as literature-anchored approximations intended to support cross-referencing rather than definitive neuroanatomical boundaries.

**Table II:**
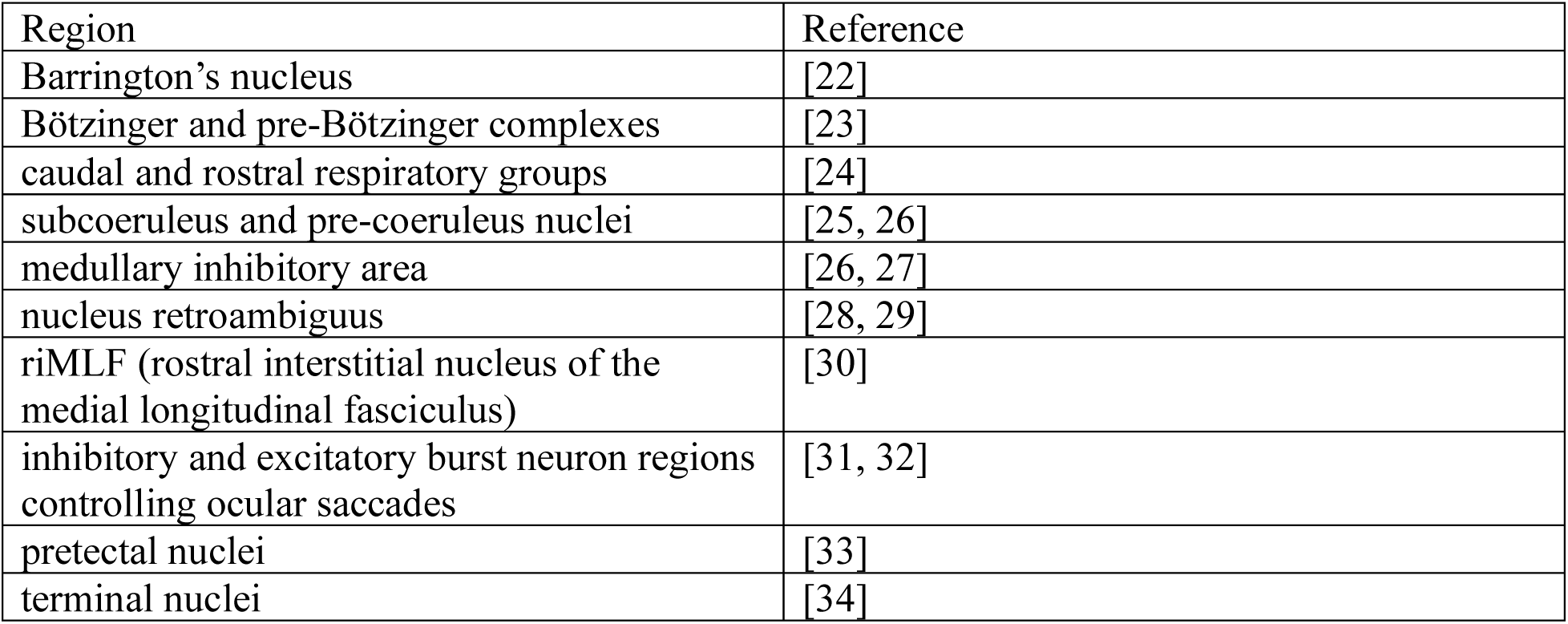
Brainstem field added to the atlas from the literature.

### Encyclopedia entry structure

Each BEAP cortical field was linked to structured encyclopedia entry synthesizing evidence from the literature. Entries included the following section (Fig. 5):

### Description

The first 1–2 sentences provide a concise summary of the main function(s) of the region, with references to review articles, meta-analyses, and/or key primary studies that clearly illustrate these functions. Subsequent paragraphs describe, in order: (i) the afferent inputs to the region (including the type of information they likely convey), (ii) how the region appears to process or transform this information, and (iii) the principal efferent targets.

This description reflects the author’s interpretation of the available data. For some controversial regions, alternative interpretations are possible and may be favored by other authors. Readers are therefore encouraged to treat this section as a guided introduction rather than a definitive statement, to consult the primary literature, and to contribute their own perspectives in the comments section.

### Alternative names

This subsection lists alternative labels used for the region of interest. These may include commonly used subregions, larger territories that encompass the region, or functional complexes. Commonly used acronyms are included alongside the full names.

### Correspondence with other atlases

Here, putative homologues of the region are listed across multiple atlases. These include macaque atlases such as the Pandya atlas [12] and the M132 atlas [4], as well as human atlases such as MMP1 and early cytoarchitectonic atlases [13-16]. Next to these atlases is an “eye” icon that, when clicked, displays the full atlas for visual comparison.

### Associated pathologies

This section lists behavioral and clinical correlates associated with lesions or atypical development involving the region of interest. It includes both individual case reports and lesion-overlap findings from larger cohorts, as well as conditions related to abnormal development (e.g., autism, schizophrenia). Each medical condition is accompanied by a brief, lay-language description of its key symptoms.

### Internal organization

This subsection summarizes studies that report functional subdivisions within the region. These studies did not meet the full criteria for defining a separate cortical field (as specified above) but nonetheless provide evidence for internal functional differentiation. Topographic maps and gradients such as somatotopy, retinotopy, and tonotopy are also described here when applicable.

### Notes

The Notes section captures relevant information that does not fit neatly into the other categories. Examples include perceptual experiences elicited by electrical stimulation, speculative or evolutionary accounts of the region’s emergence, and studies that appear to contradict or qualify the dominant functional interpretation presented in the Description.

### Border demarcation

This section documents studies, preferably fMRI-based, that clearly demarcate each of the boundaries of the region of interest and the contrasts used to reveal it. The goal is to provide practical guidance for researchers seeking to replicate or refine the boundaries used in this atlas. Border information derived from cytoarchitectonic, or molecular (e.g., genetic marker) mapping is also included here.

### Connectivity

Connectivity is organized into several subsections. First, standardized target systems that are relevant for nearly all cortical regions are summarized (e.g., dorsal thalamus, striatum, basis pontis/pontine nuclei, claustrum). Additional subsections list region-specific afferent and efferent connections, highlighting inputs and outputs that are distinctive for the cortical field in question.

### Peer review and comments

Because it is beyond the scope of a single publication to fully vet every encyclopedia entry, this responsibility is shared with the broader community of users. Readers are invited to comment on each entry, suggest corrections or improvements, and highlight additional relevant references. Unlike static journal articles, this dynamic comment system allows entries to evolve incrementally as new data and perspectives emerge.

### Platform implementation and interoperability

BEAP is publicly accessible through an interactive three-dimensional atlas interface (https://brainatlas.online/3d-brain/) and a linked encyclopedia platform (https://brainatlas.online/encyc/). Users can visualize cortical and subcortical structures, explore putative connectivity relationships, and access associated encyclopedia entries (Fig. 4-5).

**Figure 4.**
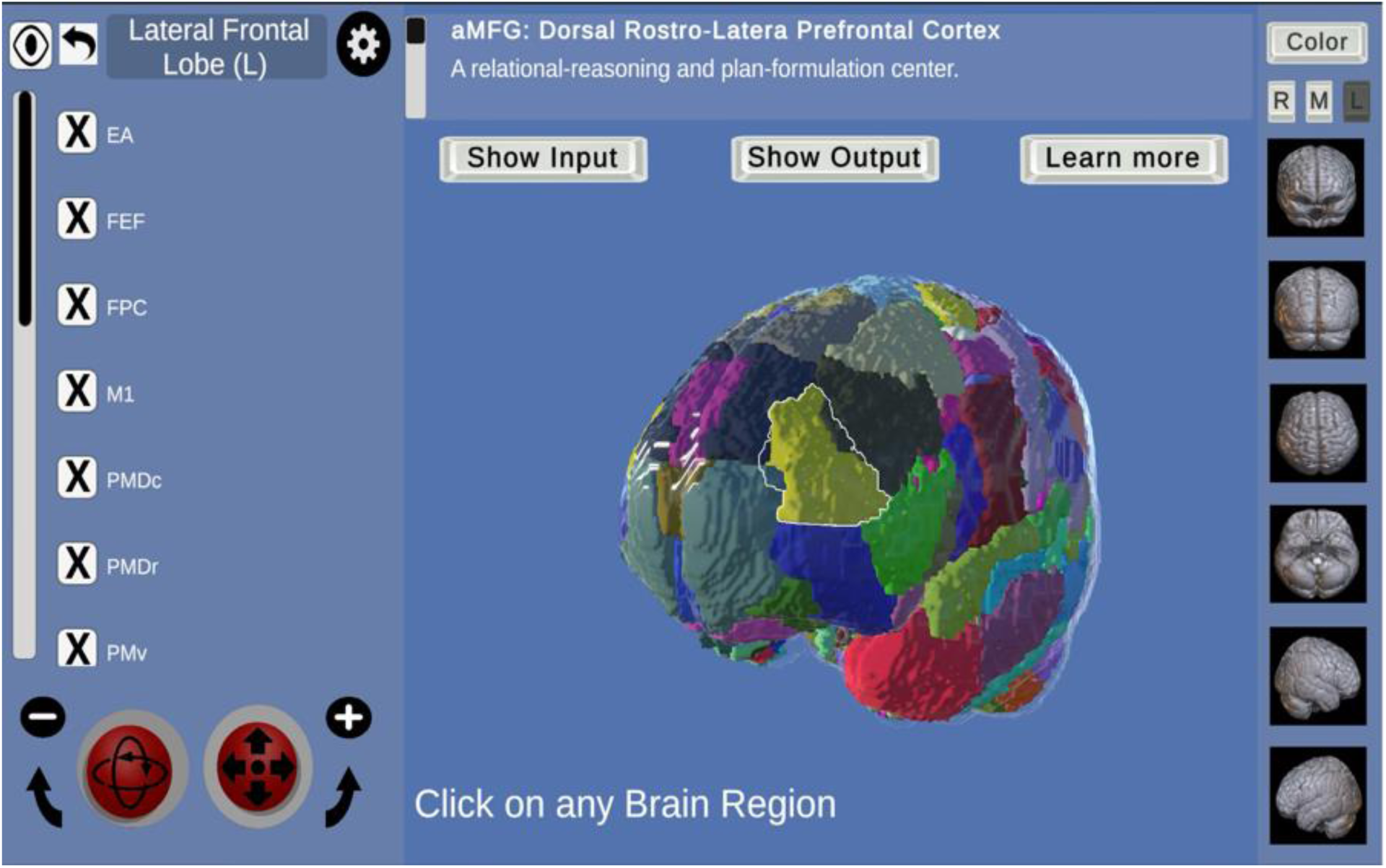
Example of the interactive three-dimensional BEAP atlas. Users can navigate the brain using the mouse or a virtual joystick (bottom-left panel). The right panel provides controls for selecting preset orientations, hiding the left, right, or midsagittal portions of the brain (buttons “R,” “M,” and “L”), and toggling color labeling to display the brain in its natural appearance. The left panel allows activation or deactivation of individual brain regions. When a region with an associated encyclopedia entry is selected, a top panel appears that allows users to display regions providing input to the selected region (afferents), regions receiving output (efferents), or to open the corresponding encyclopedia entry by clicking the “Learn more” button (Fig. 5).

**Figure 5.**
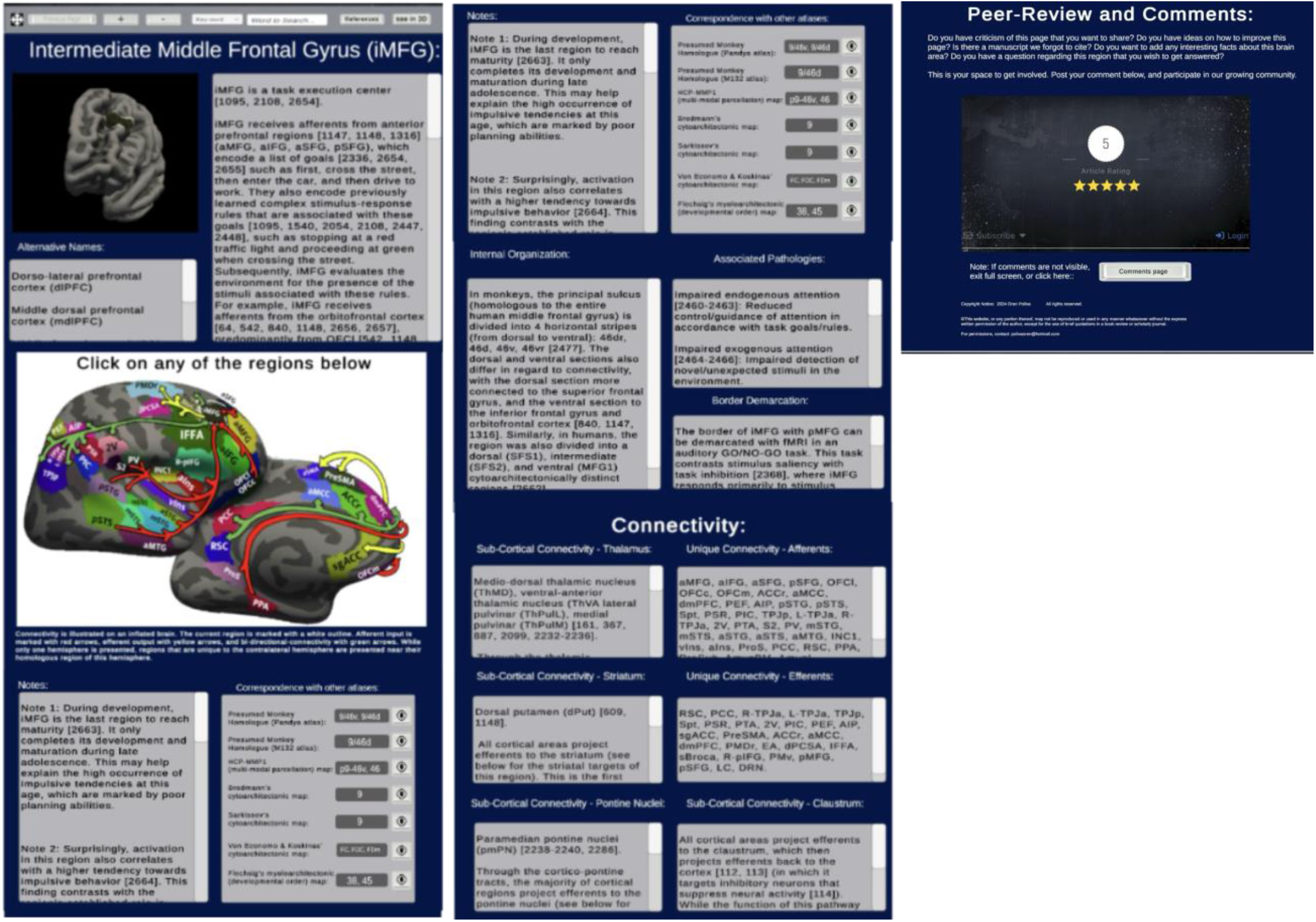
Example of a BEAP encyclopedia entry. Selecting one of the 106 cortical fields in the three-dimensional atlas (Fig. 8) opens a dedicated encyclopedia page for that region. Each page presents a representative image and a brief functional summary, followed by sections describing alternative names, correspondence with other atlases, associated pathologies, internal organization, notes, border demarcations, and lists of connections with the dorsal thalamus, striatum, basis pontis, and region-specific afferent and efferent targets. A comment section at the bottom of the page allows users to provide feedback, suggest corrections, and recommend additional references.

**Figure 6.**
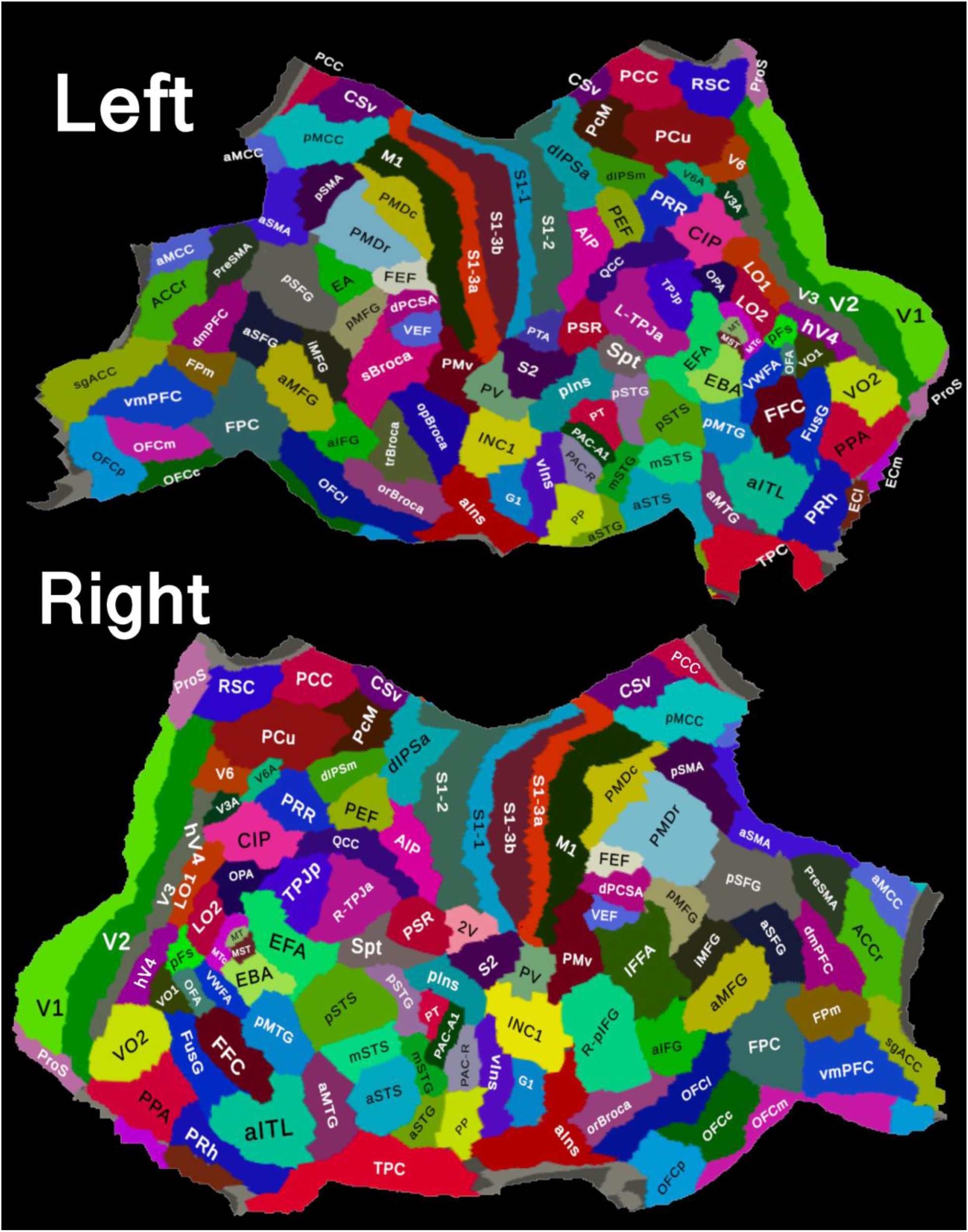
Correspondence between BEAP cortical fields and MMP1 parcels. Map showing the relationship between BEAP cortical fields and MMP1 parcels in the left hemisphere, shown here as an example.

**Figure 7.**
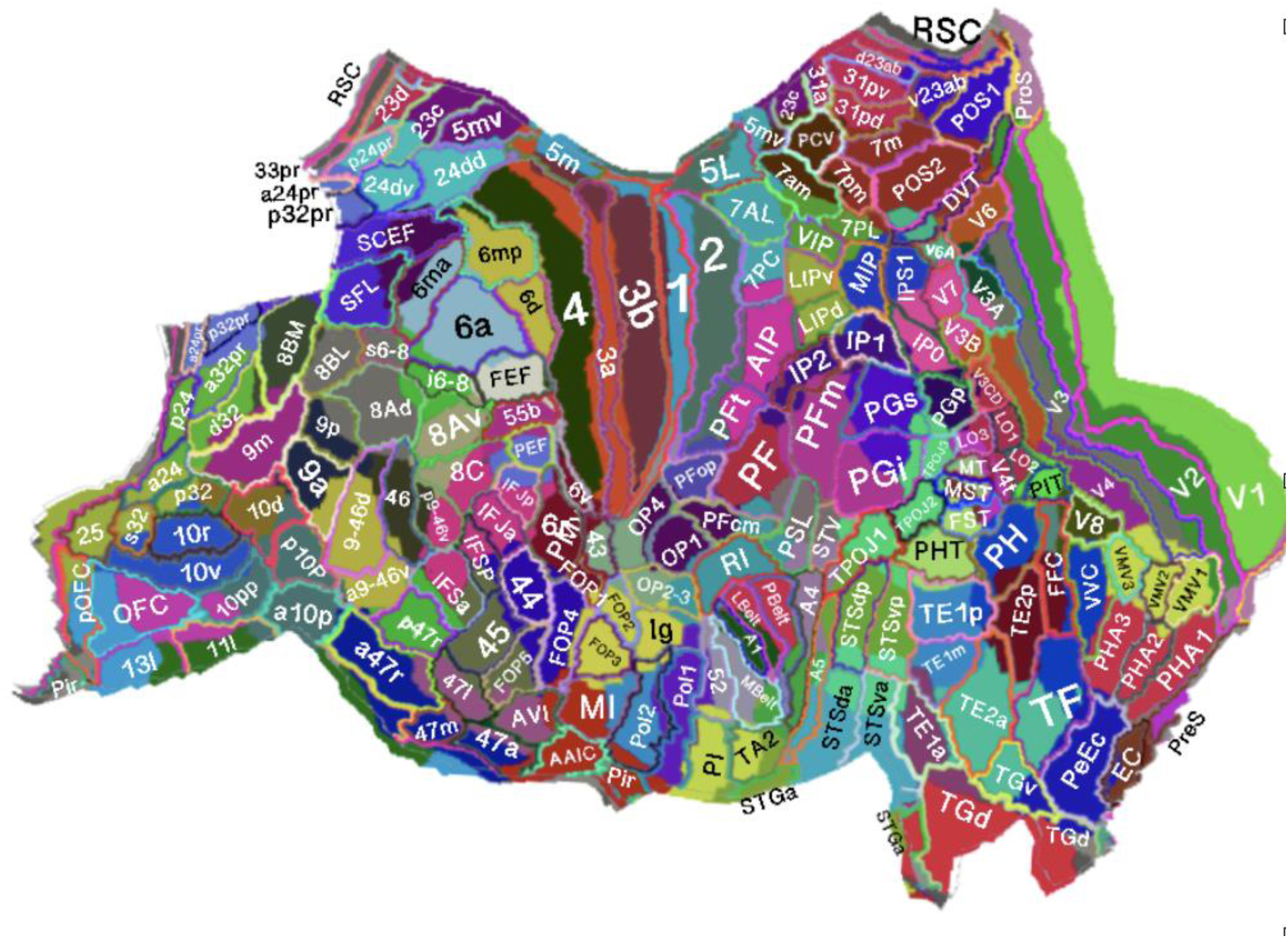
Correspondence between BEAP cortical fields and MMP1 parcels. Map showing the relationship between BEAP cortical fields (colors correspond to the fields presented in Fig. 6) and MMP1 parcels in the left hemisphere, shown here as an example.

To support interoperability with standard neuroimaging workflows, BEAP provides downloadable volumetric NIfTI files (.nii.gz), surface annotation files (.annot), and color lookup tables (.lut) compatible with software environments including FSLeyes and FreeView (https://brainatlas.online/for-researchers/). See Fig. S3 in Supplemental Material 3, for an example of integration of this atlas with FSLeyes.

### Reproducibility and transparency

To evaluate procedural stability, a subset of 100 studies (5 studies of 20 randomly selected cortices) was independently re-curated approximately one year after the initial annotation while blinded to prior assignments. Additional analyses evaluated sensitivity to the order in which studies were incorporated into the curation process.

Because BEAP is an expert-curated resource, assignments should be interpreted as literature-guided approximations subject to future refinement as additional evidence becomes available.

## RESULTS

### Contents and scope of the BEAP resource

BEAP was constructed from spatially localizable evidence extracted from 1,940 studies, comprising 1,465 human studies and 475 non-human primate tract-tracing studies. The human evidence base comprised figures from 691 fMRI, 62 PET, 44 TMS, 36 ECoG, 39 sEEG, 64 direct electrical stimulation, 16 MEG, and 513 lesion studies. Evidence relevant to border demarcation additionally included 81 post-mortem histological studies and MRI studies examining human anatomical features (e.g., cortical thickness, myelin-density imaging). Beyond figure-based localization, the BEAP resource cites 2,134 additional articles (including meta-analyses and reviews) to support functional summaries and network-level interpretations. In total, BEAP cites 3,604 manuscripts at the time of writing.

The current release of BEAP includes 340 anatomically defined regions of interest across the brain, encompassing neocortical, cerebellar, and subcortical structures. Within the neocortex, these regions are distributed throughout the occipital (20 regions), temporal (21), parietal (23) lateral frontal (21), orbitomedial frontal (14), and insular (7) lobes. Among the identified cortical fields, 98 were present bilaterally, whereas 6 were identified only in the left hemisphere and 4 only in the right hemisphere, resulting in 106 unique cortical fields across both hemispheres.

Beyond the neocortex, BEAP includes allocortical structures, comprising archi-cortical (13 regions; including hippocampal fields), and paleocortical structures (rhinencephalic/olfactory structures (6), amygdalar nuclei (11), claustrum (1), and substantia innominata (6)). Within the cerebellar region, BEAP includes cerebellar cortex (13 regions), cerebellar nuclei (7), and flocculo-nodular lobule (5). Subcortical and diencephalic structures include dorsal thalamus (28), prethalamus (7), epithalamus (1), pretectum (9), hypothalamus (16), and basal ganglia (9). Brainstem structures include midbrain (30), pons (36), and medulla (33), and the dataset further includes cervical spinal cord regions (6).

In addition, BEAP includes a set of anatomical structures displayed for visualization and spatial context, including cranial nerves (13), subcortical and cortical tracts (74), commissures (19), ventricular structures (11), and circum-ventricular organs (11). These elements are included to support anatomical orientation and navigation within the platform and were not defined through the same literature-based curation or histological criteria as the regions described above.

In the current release, encyclopedia entries were formulated only for the 106 neocortical fields and 18 cerebellar fields.

### Mapping literature-defined functional entities to a common reference framework

Using the figure-to-template localization workflow (Methods; Fig. 1), the 180 parcels per hemisphere of the MMP1 reference atlas were organized into 104 previously reported cortices in the left hemisphere and 102 in the right hemisphere. Each BEAP cortex is linked to one or more MMP1 parcels, enabling direct interoperability with standard neuroimaging workflows while preserving the literature-defined functional identity of the region. The median number of supporting functional studies was 12 (range 3–34). Entries supported by fewer than five functional studies were retained due to strong convergent non-human anatomical evidence in 2 instances (entorhinal cortices ECl and ECm) (see Methods).

Several recurring features emerged from this mapping. First, multiple BEAP cortices were associated with more than one distinguishable function within the same approximate territory, including the fusiform face complex (FFC), posterior, anterior, and ventral insular cortices (pIns, aIns, vIns), parietal ventral cortex (PV), anterior inferior temporal lobe (aITL), posterior and central orbitofrontal cortices (OFCp, OFCc), orbital part of Broca’s area (orBroca), right posterior inferior frontal gyrus (R-pIFG), anterior supplementary motor area (aSMA), and primary gustatory and viscerosensory cortex (G1), reflecting repeated reports of functional heterogeneity within these cortices.

Second, in a subset of cases, the functional profile differed across hemispheres relative to the nominal left–right homologue. Examples include the following hemispheric contrasts (right / left): second vestibular cortex (2 V) / parietal tool area (PTA), right posterior inferior frontal gyrus (R-pIFG) / opercular part of Broca’s area (opBroca)+ triangular part of Broca’s area (trBroca), inferior frontal face area (IFFA) / superior Broca’s area (sBroca), right anterior temporal-parietal junction (R-TPJa) / left anterior temporal-parietal junction (L-TPJa). These asymmetries reflect both known hemispheric lateralization and differences in how functions have been described across the literature.

Third, mapping literature-defined functional cortices to the MMP1 atlas revealed that many BEAP cortices correspond closely to existing parcel boundaries. However, in several regions the literature suggested functional distinctions that did not align fully with the MMP1 segmentation, including territories within auditory cortex, orbitofrontal cortex, insula, fusiform cortex, supplementary motor area, and intraparietal cortex (see Discussion for details).

Finally, several cortical fields identified during curation lacked standardized nomenclature across the literature despite recurring spatial and functional descriptions. In these cases, BEAP assigned stable labels while preserving alternative names and citation provenance within the corresponding encyclopedia entries. These cortices include the quantity comparison cortex (QCC), extrastriate face area (EFA), parietal tool area (PTA), posterior and anterior superior frontal gyrus (pSFG, aSFG), and ventral eye field (VEF). Unique labels were also given to sub-regions of many cortices.

### Putative connectivity integration

BEAP incorporated putative connectivity annotations derived from 475 non-human primate tract-tracing studies. Candidate macaque homologues were assigned to human cortical fields through comparison with established macaque atlases and published homology relationships. Connectivity annotations included putative afferent and efferent relationships together with standardized connectivity targets, including thalamic nuclei, striatal territories, claustrum, and pontine nuclei. These connectivity annotations were integrated directly into encyclopedia entries and interactive atlas functions, enabling users to visualize putative input and output relationships between cortical systems within the platform.

### Reproducibility and robustness of the curation procedure

Because BEAP is an expert-curated resource, the reliability of the curation framework represents an important consideration. To evaluate procedural stability, a subset of 100 studies was independently re-curated approximately one year after the initial annotation while blinded to prior assignments. While most studies (94/100) replicated the original parcel assignment, the few discrepancies involved assignment to neighboring parcels rather than non-neighboring parcels.

Additional analyses examined sensitivity to the order in which studies were incorporated into the curation process. Across representative cortical fields, the principal parcel assignments remained stable across randomized incorporation orders, with variability largely restricted to marginal parcels supported by limited evidence.

Together, these analyses suggest that the principal organizational features of the atlas are robust to reasonable variation in curation order and parcel assignment decisions. Nevertheless, because BEAP relies on expert-guided literature synthesis, some assignments necessarily involve interpretive judgment. To mitigate this limitation, the platform was designed as a transparent and versioned resource in which supporting evidence, alternative interpretations, and community feedback remain openly inspectable and subject to ongoing refinement. Accordingly, BEAP should be viewed not as a fixed endpoint, but as an evolving synthesis framework intended to improve incrementally as additional evidence and feedback accumulate.

## DISCUSSION

The Brain Encyclopedia Atlas Project (BEAP) is a literature-derived neuroinformatics resource designed to organize spatially localizable evidence about human functional neuroanatomy within a common reference framework. In the current release, BEAP integrates evidence from 1,465 human studies and 475 non-human primate tract-tracing studies to generate an interactive atlas linked directly to structured encyclopedia entries. The resource incorporates functional neuroimaging, lesion mapping, intracranial electrophysiology, cortical stimulation, and anatomical connectivity evidence, together with downloadable atlas files compatible with commonly used neuroimaging software environments. By linking literature-defined cortical fields to a standardized anatomical framework, BEAP is intended both as an educational resource and as a practical tool for contextualizing neuroimaging findings within the broader neuroscience literature.

Unlike traditional atlases that primarily emphasize anatomy, cytoarchitecture, or MRI-derived parcellation, BEAP focuses specifically on aggregating literature-defined functional territories. The resource was therefore designed not as a replacement for existing atlas frameworks, but as a complementary layer that links atlas parcels to the experimental evidence from which many functional neuroanatomical concepts originally emerged.

### Methodological contribution and curation framework

A central contribution of BEAP is the adoption of an explicit and conservative framework for synthesizing heterogeneous localization evidence across modalities. Rather than assigning cortical fields solely on the basis of isolated activations or coordinate peaks, the atlas integrates convergent evidence from functional imaging, lesion studies, intracranial recordings, cortical stimulation, and boundary-defining contrasts.

This approach reflects the empirical reality that many studies lack the spatial precision required for definitive single-parcel attribution and that cortical boundaries are often described inconsistently across experimental paradigms and modalities. BEAP therefore treats cortical fields as literature-defined entities whose spatial characterization emerges gradually as evidence accumulates across studies.

The curation framework also distinguishes between cortical fields and finer-grained internal organization. When studies suggested functional subdivisions within a territory but did not provide sufficient evidence for a distinct cortical field, these patterns were documented as sub-regions rather than used to generate new cortices. This strategy was intended to reduce premature fragmentation while preserving potentially important subregional information.

More broadly, the present framework avoids imposing artificially precise boundaries in regions where the literature provides inconsistent, overlapping, or context-dependent localization evidence. In several association cortices, published borders varied across studies or appeared graded rather than sharply delineated. In such cases, BEAP preserves these ambiguities through broader field definitions, internal organization descriptions, transition regions, and documentation of conflicting findings, rather than forcing definitive parcel assignments unsupported by all the available evidence.

### Relationship to existing atlas frameworks and neuroinformatics tools

BEAP was designed to complement existing atlas frameworks and neuroinformatics resources rather than replace them. In the present implementation, the MMP1 atlas was adopted as the primary spatial indexing framework because of its widespread use in contemporary neuroimaging workflows. Within BEAP, MMP1 functions primarily as a standardized coordinate framework that allows literature-defined cortical fields to be associated with parcels already commonly used in MRI analyses.

Across much of the neocortex, the literature-defined cortical fields identified in BEAP demonstrated broad correspondence with MMP1 parcel boundaries. This convergence suggests that multimodal high-quality MRI-derived parcellation and decades of functional localization research often identify similar large-scale cortical subdivisions. At the same time, several cortical territories exhibited functional distinctions that did not align completely with existing MMP1 boundaries, illustrating how different methodological approaches may emphasize different aspects of cortical organization (see Supplementary Material 4 for a detailed evaluation).

BEAP also differs substantially from coordinate-based meta-analytic platforms such as Neurosynth and NeuroQuery. Those tools aggregate coordinate-reported fMRI studies to estimate statistical associations between activation patterns and semantic terms. In contrast, BEAP performs a complementary region-centered synthesis that incorporates figure-based localization together with lesion studies, intracranial recordings, cortical stimulation findings, and tract-tracing evidence. Figure-based localization additionally preserves spatial context often absent from coordinate-only databases, including activation extent, relationships to sulcal landmarks, and contrasts used to demarcate neighboring regions.

Together, these characteristics position BEAP as a complementary resource within the broader neuroinformatics ecosystem, emphasizing literature synthesis, functional interpretation, and interoperability with established neuroimaging workflows.

### Practical value for research and education

BEAP is intended primarily as an interpretive and organizational resource. Each cortical field links directly to a structured encyclopedia entry summarizing convergent functional evidence, lesion associations, internal organization, boundary-defining contrasts, and putative connectivity relationships. This structure allows users to move directly from an observed activation or lesion location to the relevant supporting literature and to evaluate alternative interpretations when they exist.

To support research interoperability, BEAP provides downloadable volumetric NIfTI files, surface annotation files, and lookup tables compatible with standard neuroimaging environments such as FSLeyes and FreeView. These resources allow atlas regions to be queried directly within volumetric and surface-based neuroimaging workflows while preserving links to the underlying literature synthesis.

The platform additionally serves educational purposes by integrating information that is often distributed across multiple experimental traditions and historical naming systems into a unified spatial framework. This may facilitate navigation of the functional neuroanatomy literature for students, clinicians, and researchers working across different subfields of neuroscience.

### Reliability and robustness of the curation procedure

Because BEAP is an expert-curated resource, the reliability of the curation framework represents an important consideration. To evaluate procedural stability, a subset of 100 studies was independently re-curated approximately one year after the initial annotation while blinded to prior assignments. While most (94 studies) replicated the parcel assignment, the few discrepancies involved assignment to neighboring parcels rather than non-neighboring parcels.

Additional analyses examined sensitivity to the order in which studies were incorporated into the curation process. Across representative cortical fields, the principal parcel assignments remained stable across randomized incorporation orders, with variability largely restricted to marginal parcels supported by limited evidence.

Together, these analyses suggest that the principal organizational features of the atlas are robust to reasonable variation in curation order and parcel assignment decisions.

### Limitations and intended use

Several limitations inherent to the construction of BEAP should be emphasized. Spatial localization relied primarily on published figures and reported coordinates, both of which vary substantially in spatial resolution, smoothing, normalization procedures, and visualization practices across studies. Consequently, cortical field boundaries should be interpreted as approximate literature-derived estimates rather than precise anatomical borders.

The atlas also reflects biases in the available neuroscience literature. Functional systems such as vision, audition, language, and motor control are represented more extensively than other domains, and lesion evidence remains unevenly distributed across cortical territories. Accordingly, study density within BEAP reflects the current distribution of evidence within the literature rather than the relative biological importance of different systems.

In addition, literature coverage is necessarily incomplete and subject to continuing revision. Although explicit curation rules were used to promote internal consistency, alternative interpretations and boundary definitions are often possible, particularly within association cortex. BEAP should therefore be regarded as an evolving interpretive framework intended to support contextualization and literature synthesis rather than as a definitive cortical parcellation.

Researchers are encouraged to combine BEAP with subject-specific analyses, task-specific localizers, quantitative neuroimaging methods, and complementary atlas frameworks when defining regions of interest or interpreting functional neuroimaging findings.

### Roadmap and versioning

BEAP was designed as a living neuroinformatics resource intended to evolve as additional evidence becomes available. The online platform supports versioned releases with stable region identifiers to facilitate citability and reproducibility across studies. Future updates will incorporate newly published findings, refine boundary annotations where evidence accumulates, and expand coverage of subcortical and brainstem structures.

Each encyclopedia entry additionally includes a public comment interface that allows researchers to suggest corrections, propose alternative interpretations, and contribute additional references. Through this combination of versioned updates and community feedback, BEAP aims to remain both stable for research use and responsive to the continuing development of the neuroscience literature.

## Supporting information

5 supplemental sections

## DATA AVAILABILITY

BEAP atlas files, lookup tables, and encyclopedia entries are publicly available at: https://brainatlas.online/for-researchers/. 3D brain model available at https://brainatlas.online/3d-brain/. Neocortical map available at https://brainatlas.online/3d-brain/. Cerebellar map available at https://brainatlas.online/encyc-Cb/.

## Authors’ Contributions

OP is the sole author of this manuscript. He conceived the project framework, designed and executed the literature-synthesis and curation methodology, built the interactive three-dimensional atlas and online encyclopedia platform performed the reproducibility analyses, and drafted and revised the manuscript text.

## Acknowledgements

We thank the research assistants who supported to formulation of this resource through proofreading: Kelly Sugai, Lucas Brizolara, Anish Marrpati, Maya Labert, Evelyn Yulianto, Dylan Yu, Bailey Johnson, Fiona McNabb, Jayne Delgado.

## References

1. Eickhoff, S.B., B.T. Yeo, and S. Genon, Imaging-based parcellations of the human brain. Nature Reviews Neuroscience, 2018. 19(11): p. 672–686.

2. Glasser, M.F., et al., A multi-modal parcellation of human cerebral cortex. Nature, 2016. 536(7615): p. 171–178.

3. Felleman, D.J. and D.C. Van Essen, Distributed hierarchical processing in the primate cerebral cortex. Cerebral cortex (New York, NY: 1991), 1991. 1(1): p. 1–47.

4. Markov, N.T., et al., A weighted and directed interareal connectivity matrix for macaque cerebral cortex. Cerebral cortex, 2014. 24(1): p. 17–36.

5. Friston, K.J., Functional and effective connectivity: a review. Brain connectivity, 2011. 1(1): p. 13–36.

6. Van Dijk, K.R., et al., Intrinsic functional connectivity as a tool for human connectomics: theory, properties, and optimization. Journal of neurophysiology, 2010. 103(1): p. 297–321.

7. Sultan, F. and M. Glickstein, The cerebellum: comparative and animal studies. The cerebellum, 2007. 6(3): p. 168–176.

8. Mai, J.K., M. Majtanik, and G. Paxinos, Atlas of the human brain. 2015: Academic press.

9. Ding, S.L., et al., Comprehensive cellular-resolution atlas of the adult human brain. Journal of comparative neurology, 2016. 524(16): p. 3127–3481.

10. Mai, J.K. and G. Paxinos, The human nervous system. 2011: Academic press.

11. Amunts, K., et al., BigBrain: an ultrahigh-resolution 3D human brain model. science, 2013. 340(6139): p. 1472–1475.

12. Yeterian, E.H., et al., The cortical connectivity of the prefrontal cortex in the monkey brain. Cortex, 2012. 48(1): p. 58–81.

13. Brodmann, K., Vergleichende lokalisationslehre der grobhirnrinde, in Vergleichende lokalisationslehre der grobhirnrinde. 1909. p. 324–324.

14. von Economo, C.F. and G.N. Koskinas, Die cytoarchitektonik der hirnrinde des erwachsenen menschen. 1925: J. Springer.

15. Sarkissov, S.A., et al., Atlas of the Cytoarchitectonics of the Human Cerebral Cortex. Medgiz. Moscow., 1955.

16. Flechsig, P.E., Anatomie des menschlichen Gehirns und Rückenmarks auf myelogenetischer Grundlage. Vol. 1. 1920: G. Thieme.

17. Seubert, J., et al., Statistical localization of human olfactory cortex. Neuroimage, 2013. 66: p. 333–342.

18. Kedo, O., et al., Receptor-driven, multimodal mapping of the human amygdala. Brain Structure and Function, 2018. 223(4): p. 1637–1666.

19. Schultz, H., T. Sommer, and J. Peters, Direct evidence for domain-sensitive functional subregions in human entorhinal cortex. Journal of Neuroscience, 2012. 32(14): p. 4716–4723.

20. Palomero-Gallagher, N., et al., Multimodal mapping and analysis of the cyto-and receptorarchitecture of the human hippocampus. Brain Structure and Function, 2020. 225: p. 881–907.

21. Averbeck, B.B., et al., Estimates of projection overlap and zones of convergence within frontal-striatal circuits. Journal of Neuroscience, 2014. 34(29): p. 9497–9505.

22. Blanco, L., et al., Critical evaluation of the anatomical location of the Barrington nucleus: relevance for deep brain stimulation surgery of pedunculopontine tegmental nucleus. Neuroscience, 2013. 247: p. 351–363.

23. Schwarzacher, S.W., U. Rüb, and T. Deller, Neuroanatomical characteristics of the human pre-Bötzinger complex and its involvement in neurodegenerative brainstem diseases. Brain, 2011. 134(1): p. 24–35.

24. Ellenberger, H. and J. Feldman, Brainstem connections of the rostral ventral respiratory group of the rat. Brain research, 1990. 513(1): p. 35–42.

25. Lu, J., et al., A putative flip–flop switch for control of REM sleep. Nature, 2006. 441(7093): p. 589–594.

26. Peever, J. and P.M. Fuller, The biology of REM sleep. Current biology, 2017. 27(22): p. R1237–R1248.

27. Karlsson, K. and M. Blumberg, Active medullary control of atonia in week-old rats. Neuroscience, 2005. 130(1): p. 275–283.

28. Aydogdu, I., et al., Dysphagia in lateral medullary infarction (wallenberg’s syndrome) an acute disconnection syndrome in premotor neurons related to swallowing activity? Stroke, 2001. 32(9): p. 2081–2087.

29. VanderHorst, V., E. Terasawa, and H. Ralston III, Monosynaptic projections from the nucleus retroambiguus region to laryngeal motoneurons in the rhesus monkey. Neuroscience, 2001. 107(1): p. 117–125.

30. Horn, A.K. and J.A. Büttner-Ennever, Brainstem circuits controlling lid–eye coordination in monkey. Progress in Brain Research, 2008. 171: p. 87–95.

31. Strassman, A., S. Highstein, and R. McCrea, Anatomy and physiology of saccadic burst neurons in the alert squirrel monkey. I. Excitatory burst neurons. Journal of Comparative Neurology, 1986. 249(3): p. 337–357.

32. Strassman, A., S. Highstein, and R. McCrea, Anatomy and physiology of saccadic burst neurons in the alert squirrel monkey. II. Inhibitory burst neurons. Journal of Comparative Neurology, 1986. 249(3): p. 358–380.

33. Hutchins, B. and J.T. Weber, The pretectal complex of the monkey: a reinvestigation of the morphology and retinal terminations. Journal of Comparative Neurology, 1985. 232(4): p. 425–442.

34. Cooper, H. and M. Magnin, Accessory optic system of an anthropoid primate, the gibbon (Hylobates concolor): evidence of a direct retinal input to the medial terminal nucleus. Journal of Comparative Neurology, 1987. 259(4): p. 467–482.

35. Chao, L.L., J. Weisberg, and A. Martin, Experience-dependent modulation of category-related cortical activity. Cerebral Cortex, 2002. 12(5): p. 545–551.

36. Yen, C.-P., et al., Impact of bilateral anterior cingulotomy on neurocognitive function in patients with intractable pain. Journal of Clinical Neuroscience, 2009. 16(2): p. 214–219.

37. Singer, T., et al., Empathy for pain involves the affective but not sensory components of pain. Science, 2004. 303(5661): p. 1157–1162.

