## Supplementary material for "The Brain Encyclopedia Atlas Project (BEAP): A Literature-Synthesis-Derived Functional Atlas of the Human Brain": 5 supplemental sections: SupplementaryMaterial2.pdf

**Somatosensory cortex:** The designation of area 3b and 3a as primary sensory cortex were known since the early maps of Brodmann and Vogt [1, 2]. Interestingly, the region medial to 3a and 3b, which Brodmann designated as area 5, was shown later, through the usage of cortical stimulation by Penfield, to encode the pelvis, including genitalia, and lower limb somatic representations [3]. Similarly, using relative myelin density MRI protocol, the medial region 5m was recently reported with the same myelin density as areas 3a and 3b [4]. In accordance with these findings, the border of S1-3a and S1-3b of the current resource is not limited to areas 3a and 3b of Brodman and Vogt, and additionally constitute the rostral and caudal sections of 5m respectively. Moreover, based on the cytoarchitectonic atlas of von Economo and Koskinas [5], which designated their area PA1 (comparable with 3a) to stretch caudally at its most medial location and directly connect with medial area PA2 (comparable to Brodmann area 2), the medial border of S1-3a of the current atlas was manually redrawn to connect with medial S1-2 (see figure S1 in supplemental material).

**Posterior/middle dorsal insula (VS, INC1, PIVC, G1):** The granular and dysgranular insula are located ventrally to the primary somatosensory cortex and similarly participate in encoding somatosensations. Like 3a and 3b, these regions also receive ascending afferents from the same thalamic complex, the ventro-lateral thalamic nuclear group [6-11].

**VS+INC1**: The posterior half of the posterior insula receives input from the posterior ventromedial thalamic nucleus (ThVMpo), which is the target of the spino-thalamic tract (the pathway encoding the protopathic sensations). An interesting finding of the current analysis is a division of labor in this territory with a posterior section, encoding the protopathic sensations of temperature, itch, and pleasant light touch (presented here with the label ventral somatosensory cortex, VS, based on previous nomenclature in monkeys [12]), and with an anterior section, encoding chiefly nociception (presented here with the new label, INC1). The borders between these regions is identifiable when the area is investigated using electrical stimulation, with stimulation to area VS reported to induce temperature sensations and INC1 pain [13].

**PIVC**: The posterior insula additionally includes the primary vestibular cortex, PIVC. This region receives ascending afferents from a different thalamic nucleus than VS and INC1, the medial pulvinar and superior ventro-posterior nuclei (VPs, ThPulM) [14, 15], which receives in turn afferents from the vestibular complex of the pons. The location of PIVC in this resource is based on a study combining cytoarchitectonic and fMRI evidence [16]. However, to what degree this region is distinct from its neighbors VS and INC1 (if indeed distinct) still needs to be established.

**G1**: In contrast to S1-3a and S1-3b, which encode the sensations from the exteroceptive surface (e.g., skin, lips, etc), the dysgranular insula shows a transition towards inner somatic representations, including the heart, lungs, digestive tract, and taste sensations from the tongue. Interestingly, in monkeys only a small subsection of this region (only 3% of the neurons [17, 18]) actually encode taste with the remaining neurons encoding viscerosenations. Based on this finding, the area was deemed in the current resource to contain two distinct regions a primary cortex for taste, G1, and a primary cortex for visceral sensations, Visc1 (the term G1, however, was used in the current resource as it is more commonly used). The two cortices are not completely unrelated as the topography here appears imported from the topographic map of the solitary nucleus, in which the rostral section encodes taste, middle section encodes the digestive tract, and the caudal section the heart and lungs [19].

**Auditory cortex (PAC-R, PAC-A1):** Historically, Heschl's gyrus was regarded as the primary auditory cortex, and is known as A1. However, following tonotopic studies in monkeys, two additional primary auditory fields, each with its own tonotopic map was reported in moneys, rostral area R, and rostrotemporal area RT [20, 21]. Area R (labelled PAC-R in the current resource) was further confirmed in humans [22-32]. Functionally. PAC-R differs from PAC-A1 for its further involvement in sound recognition [33-35].

**PT**: Interestingly, the area posterior to PAC-A1, the planum temporale (PT in the current resource; aka area Tpt or areas CM and CL), is regarded as a secondary (associative) auditory cortex [36]. This chiefly the outcome of a monkey study by Rauschecker et al [37], who reported that following lesions of A1, 70-80% of the neurons in CM lost their responsivity to sounds. However, the authors do acknowledge that the remaining neurons still responded to sounds (although the response was to complex sounds, not pure tones). Nonetheless, later studies showed that neurons here respond to sound (mostly complex sounds) before any response is registered in areas A1 or R [38, 39]. Recently, using intra-cranial recordings similar findings were reported in humans [40]. This implies that not only the posterior auditory cortex is a primary auditory cortex, it further indicates of an alternative sub-cortical auditory processing stream, in which complex sounds are analyzed and recognized independently to pure tones.

**ProS:** Located anterior to V1, anterior to the section of V1 encoding the extreme visual periphery, there is an additional retinotopic field, known as area ProS. Remarkably, a large percentage of neurons here become responsive to flashes of light before responsiveness is recorded in most V1 neurons (at approximately 35 ms latency) [41]. Functionally, ProS appears specialized for the detection of moving objects at the visual periphery [41-43], and thus having this region receiving direct ascending afferent input, bypassing V1, provides an adaptive advantage.

**MT**: The visual cortex responsible for the detection of localized coherent visual motion, the middle temporal cortex, also exhibits the characteristics of a primary sensory cortex. First, it receives direct ascending afferent input from the retina via relay stations in the superior colliculus and inferior pulvinar thalamic nucleus (ThPuli) [44, 45]. Using both cytoarchitectonic mapping and relative myelin density MRI [4, 46], MT was also was shown with a very dense underlying myelin sheet, which is consistent with primary sensory regions. Moreover, in monkeys, a small percentage of MT neurons showed responsiveness to visual stimuli before V1 [47]. Using MEG similar latency for the two regions was also reported in humans [48]. Finally, in the classic study of Flechsig, which mapped cortical fields in post-mortem human fetuses based on myeloarchitectonic developmental order, MT was shown myelinated during the fetal pre-natal period [49] (like other sensory cortices).

**PEF**: The most counter-intuitive inclusion into this category is PEF. However, like other sensory cortices, PEF exhibits a dense underlying myelin sheet [4, 50, 51]. Moreover, like MT and other sensory cortices, Flechsig reported this region to develop prenatally [49]. This is in contrast to all its parietal neighbors which become myelinated during the first year of life. Finally, recordings in both monkeys and humans showed this region to respond to visual stimuli with similar latency to V1 [52, 53], and di-synaptic labelling in monkeys showed this region to receive afferent input that originate in the superior colliculus [54].


1. Brodmann, K., *Vergleichende lokalisationslehre der grobhirnrinde*, in *Vergleichende lokalisationslehre der grobhirnrinde*. 1909. p. 324-324.
2. Vogt, C. and O. Vogt, *Allgemeine ergebnisse unserer hirnforschung*. Vol. 25. 1919: JA Barth.
3. Penfield, W. and T. Rasmussen, *The cerebral cortex of man; a clinical study of localization of function.* 1950.

4. Glasser, M.F. and D.C. Van Essen, *Mapping human cortical areas in vivo based on myelin content as revealed by T1-and T2-weighted MRI.* Journal of neuroscience, 2011. **31**(32): p. 11597-11616.
5. von Economo, C.F. and G.N. Koskinas, *Die cytoarchitektonik der hirnrinde des erwachsenen menschen*. 1925: J. Springer.
6. Craig, A., *Topographically organized projection to posterior insular cortex from the posterior portion of the ventral medial nucleus in the long-tailed macaque monkey.* Journal of Comparative Neurology, 2014. **522**(1): p. 36-63.
7. Krause, T., et al., *Thalamic sensory strokes with and without pain: differences in lesion patterns in the ventral posterior thalamus.* Journal of Neurology, Neurosurgery & Psychiatry, 2012. **83**(8): p. 776-784.
8. Lenz, F., *The ventral posterior nucleus of thalamus is involved in the generation of central pain syndromes.* APS Journal, 1992. **1**(1): p. 42-51.
9. Dum, R.P., D.J. Levinthal, and P.L. Strick, *The spinothalamic system targets motor and sensory areas in the cerebral cortex of monkeys.* Journal of Neuroscience, 2009. **29**(45): p. 14223-14235.
10. Blomquist, A.J., R.M. Benjamin, and R. Emmers, *Thalamic localization of afferents from the tongue in squirrel monkey (Saimiri sciureus).* Journal of Comparative Neurology, 1962. **118**(1): p. 77-87.
11. Pritchard, T., et al., *Projections of thalamic gustatory and lingual areas in the monkey, Macaca fascicularis.* Journal of Comparative Neurology, 1986. **244**(2): p. 213-228.
12. Cusick, C., et al., *Somatotopic organization of the lateral sulcus of owl monkeys: area 3b, S-II, and a ventral somatosensory area.* Journal of Comparative Neurology, 1989. **282**(2): p. 169-190.
13. Afif, A., et al., *Anatomofunctional organization of the insular cortex: a study using intracerebral electrical stimulation in epileptic patients.* Epilepsia, 2010. **51**(11): p. 2305-2315.
14. Akbarian, S., O.J. Grüsser, and W. Guldin, *Thalamic connections of the vestibular cortical fields in the squirrel monkey (Saimiri sciureus).* Journal of Comparative Neurology, 1992. **326**(3): p. 423-441.
15. Lopez, C. and O. Blanke, *The thalamocortical vestibular system in animals and humans.* Brain Res Rev, 2011. **67**(1-2): p. 119-46.
16. Eickhoff, S.B., et al., *Identifying human parieto-insular vestibular cortex using fMRI and cytoarchitectonic mapping.* Human brain mapping, 2006. **27**(7): p. 611-621.
17. Scott, T.R., et al., *Gustatory neural coding in the monkey cortex: stimulus intensity.* Journal of Neurophysiology, 1991. **65**(1): p. 76-86.
18. Smith, D.V. and S.J. St John, *Neural coding of gustatory information.* Current opinion in neurobiology, 1999. **9**(4): p. 427-435.
19. Cutsforth-Gregory, J.K. and E.E. Benarroch, *Nucleus of the solitary tract, medullary reflexes, and clinical implications.* Neurology, 2017. **88**(12): p. 1187-1196.
20. Kaas, J.H. and T.A. Hackett, *Subdivisions of auditory cortex and processing streams in primates.* Proceedings of the National Academy of Sciences, 2000. **97**(22): p. 11793-11799.

21. Tian, B., et al., *Functional specialization in rhesus monkey auditory cortex.* Science, 2001. **292**(5515): p. 290-293.
22. Saenz, M. and D.R. Langers, *Tonotopic mapping of human auditory cortex.* Hearing research, 2014. **307**: p. 42-52.
23. Langers, D.R., *Assessment of tonotopically organised subdivisions in human auditory cortex using volumetric and surface-based cortical alignments.* Human brain mapping, 2014. **35**(4): p. 1544-1561.
24. Da Costa, S., et al., *Tonotopic gradients in human primary auditory cortex: concurring evidence from high-resolution 7 T and 3 T fMRI.* Brain Topogr, 2015. **28**(1): p. 66-9.
25. van Dijk, P. and D.R. Langers, *Mapping tonotopy in human auditory cortex*, in *Basic aspects of hearing: Physiology and perception*. 2013, Springer. p. 419-425.
26. Striem-Amit, E., U. Hertz, and A. Amedi, *Extensive cochleotopic mapping of human auditory cortical fields obtained with phase-encoding FMRI.* PloS one, 2011. **6**(3): p. e17832.
27. Moerel, M., F. De Martino, and E. Formisano, *Processing of natural sounds in human auditory cortex: tonotopy, spectral tuning, and relation to voice sensitivity.* Journal of Neuroscience, 2012. **32**(41): p. 14205-14216.
28. Dick, F., et al., *In vivo functional and myeloarchitectonic mapping of human primary auditory areas.* Journal of Neuroscience, 2012. **32**(46): p. 16095-16105.
29. Formisano, E., et al., *Mirror-symmetric tonotopic maps in human primary auditory cortex.* Neuron, 2003. **40**(4): p. 859-869.
30. Talavage, T.M., et al., *Tonotopic organization in human auditory cortex revealed by progressions of frequency sensitivity.* Journal of neurophysiology, 2004. **91**(3): p. 1282-1296.
31. Humphries, C., E. Liebenthal, and J.R. Binder, *Tonotopic organization of human auditory cortex.* Neuroimage, 2010. **50**(3): p. 1202-1211.
32. Petkov, C.I., et al., *Functional imaging reveals numerous fields in the monkey auditory cortex.* PLoS biology, 2006. **4**(7): p. e215.
33. Steinschneider, M., et al., *Intracortical responses in human and monkey primary auditory cortex support a temporal processing mechanism for encoding of the voice onset time phonetic parameter.* Cerebral Cortex, 2005. **15**(2): p. 170-186.
34. Yin, P., et al., *Early stages of melody processing: stimulus-sequence and task-dependent neuronal activity in monkey auditory cortical fields A1 and R.* Journal of neurophysiology, 2008. **100**(6): p. 3009-3029.
35. Poliva, O., et al., *Functional mapping of the human auditory cortex: fMRI investigation of a patient with auditory agnosia from trauma to the inferior colliculus.* Cognitive and Behavioral Neurology, 2015. **28**(3): p. 160-180.
36. Griffiths, T.D. and J.D. Warren, *The planum temporale as a computational hub.* Trends in neurosciences, 2002. **25**(7): p. 348-353.
37. Rauschecker, J.P., et al., *Serial and parallel processing in rhesus monkey auditory cortex.* Journal of Comparative Neurology, 1997. **382**(1): p. 89-103.

38. Camalier, C.R., et al., *Neural latencies across auditory cortex of macaque support a dorsal stream supramodal timing advantage in primates.* Proceedings of the National Academy of Sciences, 2012. **109**(44): p. 18168-18173.
39. Lakatos, P., et al., *Timing of pure tone and noise-evoked responses in macaque auditory cortex.* Neuroreport, 2005. **16**(9): p. 933-937.
40. Hamilton, L.S., et al., *Parallel and distributed encoding of speech across human auditory cortex.* Cell, 2021. **184**(18): p. 4626-4639. e13.
41. Yu, H.-H., et al., *A specialized area in limbic cortex for fast analysis of peripheral vision.* Current Biology, 2012. **22**(14): p. 1351-1357.
42. Palmer, S.M. and M. Rosa, *A distinct anatomical network of cortical areas for analysis of motion in far peripheral vision.* European Journal of Neuroscience, 2006. **24**(8): p. 2389-2405.
43. Tamietto, M. and D.A. Leopold, *Visual Cortex: The Eccentric Area Prostriata in the Human Brain.* Curr Biol, 2018. **28**(1): p. R17-R19.
44. Shipp, S., *The functional logic of cortico–pulvinar connections.* Philosophical Transactions of the Royal Society of London. Series B: Biological Sciences, 2003. **358**(1438): p. 1605-1624.
45. Sincich, L.C., et al., *Bypassing V1: a direct geniculate input to area MT.* Nature neuroscience, 2004. **7**(10): p. 1123-1128.
46. Tootell, R.B. and J.B. Taylor, *Anatomical evidence for MT and additional cortical visual areas in humans.* Cerebral Cortex, 1995. **5**(1): p. 39-55.
47. Schmolesky, M.T., et al., *Signal timing across the macaque visual system.* Journal of neurophysiology, 1998. **79**(6): p. 3272-3278.
48. Ffytche, D.H., C. Guy, and S. Zeki, *The parallel visual motion inputs into areas V1 and V5 of human cerebral cortex.* Brain, 1995. **118**(6): p. 1375-1394.
49. Flechsig, P.E., *Anatomie des menschlichen Gehirns und Rückenmarks auf myelogenetischer Grundlage*. Vol. 1. 1920: G. Thieme.
50. Siegel, R., et al., *The functional and anatomical subdivision of the inferior parietal labule.* Society for Neuroscience Abstracts. In press.[RAA], 1985.
51. Glasser, M.F., et al., *A multi-modal parcellation of human cerebral cortex.* Nature, 2016. **536**(7615): p. 171-178.
52. Regev, T.I., et al., *Human posterior parietal cortex responds to visual stimuli as early as peristriate occipital cortex.* European Journal of Neuroscience, 2018. **48**(12): p. 3567-3582.
53. Bisley, J.W., B.S. Krishna, and M.E. Goldberg, *A rapid and precise on-response in posterior parietal cortex.* Journal of Neuroscience, 2004. **24**(8): p. 1833-1838.
54. Ugolini, G. and W. Graf, *Pathways from the superior colliculus and the nucleus of the optic tract to the posterior parietal cortex in macaque monkeys: Functional frameworks for representation updating and online movement guidance.* European Journal of Neuroscience, 2024.
