## Supplementary material for "The Brain Encyclopedia Atlas Project (BEAP): A Literature-Synthesis-Derived Functional Atlas of the Human Brain": 5 supplemental sections: SupplementaryMaterial3.pdf

Supplementary Material 3:

Additional Figures:

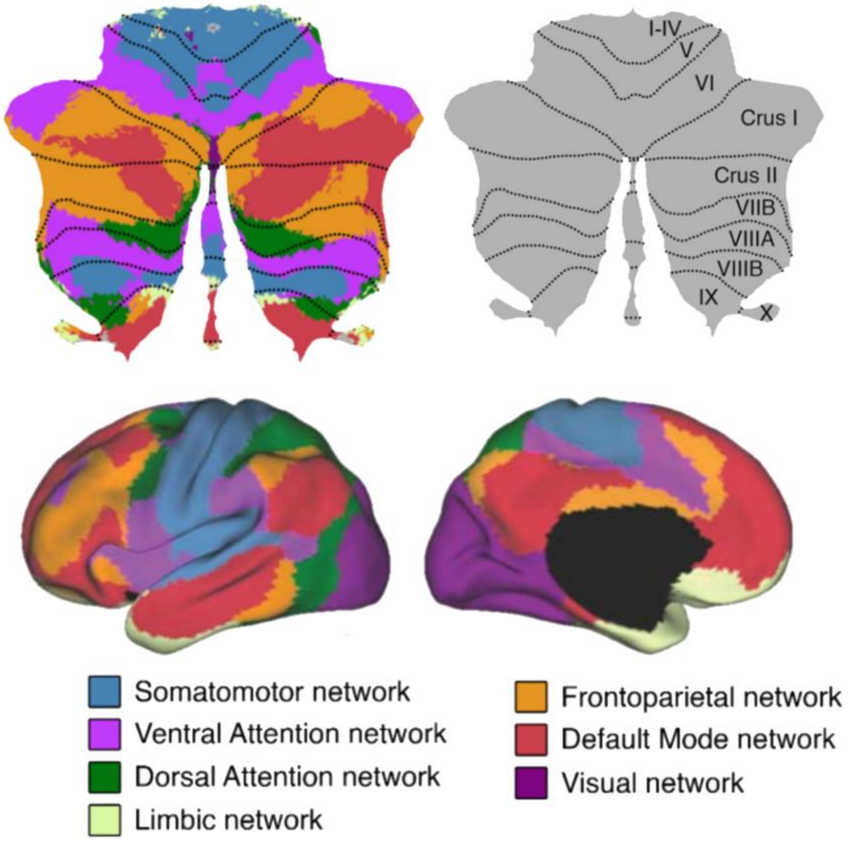


**Figure S1. Mapping of the cerebellar fields.**
Using resting-state fMRI, Buckner et al. [21] mapped the spatial distribution of seven large-scale functional networks across the cerebellar cortex by correlating cerebellar activity with co-active regions in the cerebral cortex. This organization can be interpreted as a topographic reflection of frontal lobe systems within the cerebellum. Specifically, primary motor cortex (blue) corresponds to cerebellar anterior lobule (I-IV, V) and lobule VIIIB; premotor cortex (green and purple) maps onto lobules VI and VIIA–VIIB; lateral prefrontal cortex (orange) corresponds to Crus I (superior and lateral portions) and Crus II (inferior and lateral portions); and medial prefrontal cortex (red) maps onto Crus I (inferior portion) and Crus II (superior portion). Figure was adapted and modified from [67] (eLife, CC BY 4.0).

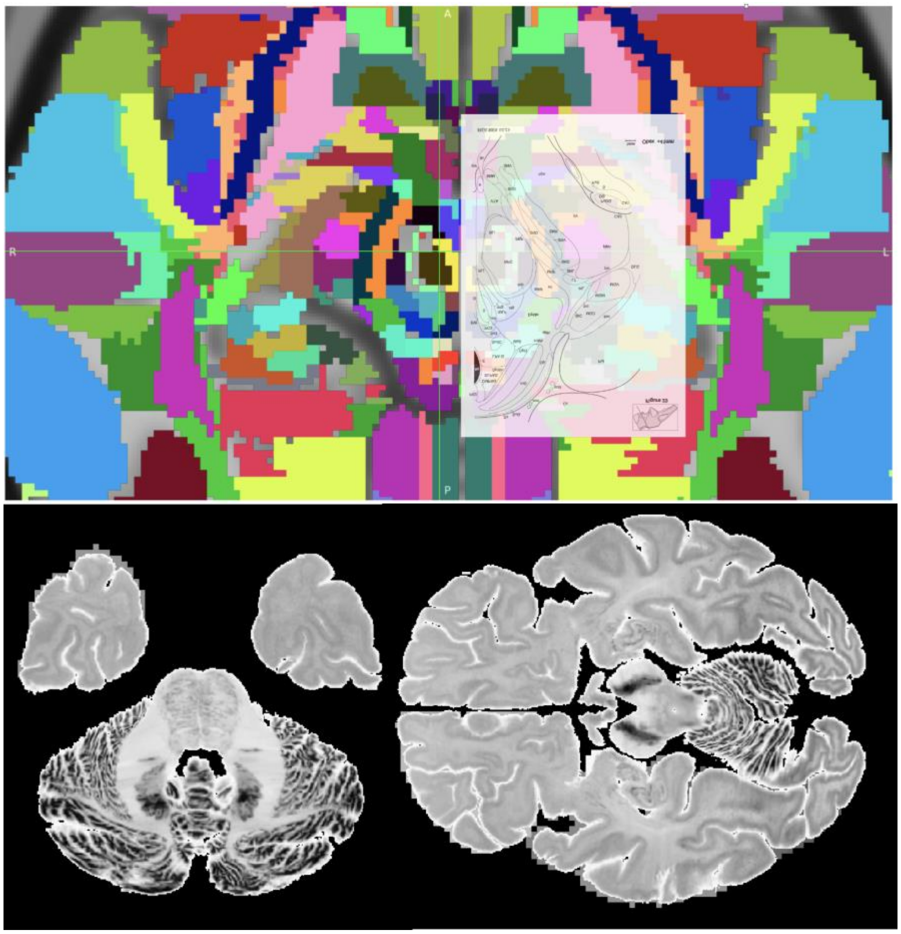

**Figure S2. Delineation of brainstem and subcortical nuclei in template space.**
Brainstem and subcortical nuclei were manually delineated on the MNI152 template using published illustrated atlases. **Top:** An axial section of the midbrain from a histological atlas was aligned to the corresponding MNI152 slice in FSLeyes and used to guide manual tracing of brainstem nuclei. **Bottom:** Several nuclei that are difficult to identify reliably on conventional MRI, including the substantia nigra, deep cerebellar nuclei, and the basis pontis (pontine nuclei), were delineated using the Big Brain Project [68], which is available in MNI152 space.
