## Supplementary material for "The Brain Encyclopedia Atlas Project (BEAP): A Literature-Synthesis-Derived Functional Atlas of the Human Brain": 5 supplemental sections: SupplementaryMaterial4.pdf

### Supplemental Material 4.

BEAP was designed to operate within an established neuroimaging reference framework. For this purpose, the Human Connectome Project multimodal parcellation atlas (HCP-MMP1) was adopted as the spatial indexing system for the present resource [1]. Using a multimodal MRI approach integrating cortical thickness, relative myelin content, resting-state connectivity, and task-fMRI data, the MMP1 atlas subdivided the neocortex into 180 parcels per hemisphere. While the delineation of parcel boundaries in MMP1 was based on highly systematic acquisition and analysis procedures, the assignment of parcel labels and their correspondence to prior neuroanatomical literature was necessarily more interpretive, often relying on a limited number of representative studies or inferred homologies (Fig. S1). The present work expands upon this process by systematically relating literature-defined cortical fields to the MMP1 framework.

Across much of the neocortex, the literature-defined cortices identified in BEAP correspond closely to parcel boundaries proposed in the MMP1 atlas, suggesting substantial convergence between multimodal MRI-based parcellation and decades of functional neuroanatomical research. At the same time, the present synthesis highlights several instances in which literature-defined functional distinctions do not map neatly onto existing parcel boundaries. These cases do not necessarily imply inaccuracies in the MMP1 atlas, but rather illustrate how different methodological approaches can emphasize partially distinct aspects of cortical organization. The examples below illustrate representative cases of convergence, divergence, and reinterpretation between literature-defined cortical fields and multimodal MRI-based parcellation schemes.

| Parcel Index | Area Name | Area Description | New? | Sections | Other Names | Key Studies |
|---|---|---|---|---|---|---|
| 44 | 6ma | Area 6m anterior | Yes* | **7**,8,22 | SMAr, 6, SMA | **Fischl et al 2008**, Vorobiev et al 1998, Geyer 2004 |

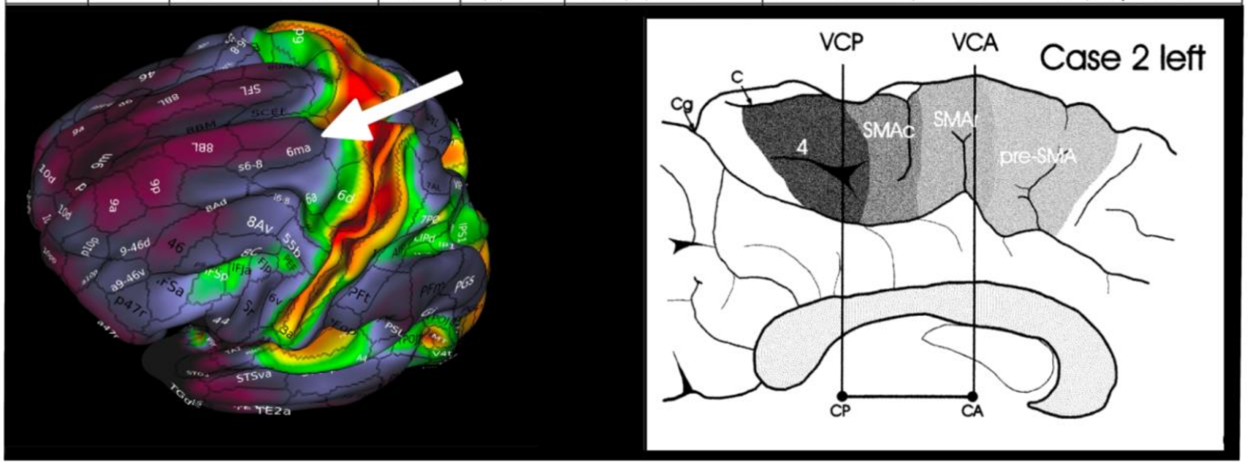


**Figure S1. Example illustrating the difficulty of linking atlas parcels to the functional neuroimaging literature.**
The Human Connectome Project multimodal parcellation atlas (MMP1) divides the neocortex into 180 parcels per hemisphere based on multimodal MRI measurements. However, assigning these parcels to previously described functional neuroanatomical entities is not always straightforward. An illustrative example is parcel 6ma, a small region located along the posterior superior frontal gyrus (left). In the supplementary material accompanying Glasser et al. (2016), parcel labels were associated with representative supporting citations. In this case, parcel 6ma was identified as part of the supplementary motor area (SMA) based on three references (top). Closer examination reveals that two of these sources [2, 3] discuss the SMA only in broad terms without explicitly defining whether it extends onto the lateral cortical surface, while the third reference [4] provides a detailed cytoarchitectonic analysis restricted primarily to the medial surface of the hemisphere (right). Consequently, the cited literature does not clearly establish whether parcel 6ma should be considered part of the SMA, illustrating the broader challenge of relating atlas parcels to explicit functional and anatomical evidence from the literature.

### Auditory Cortex

One example concerns auditory cortex. In the MMP1 atlas, associative auditory cortex along the superior temporal gyrus is subdivided into ventral-to-dorsal strips labeled A4 and A5, based largely on reductions in myelin density observed with relative myelin content imaging. The literature synthesis conducted here suggests that functional distinctions within this territory are more commonly described along a posterior-anterior axis, with the posterior superior temporal gyrus commonly associated with auditory spatial processing [5-9], and the more anteriorly located middle superior temporal gyrus with analysis of pitch features, including speaker discrimination [10, 11], speaker recognition [12-16], gender estimation [17], and extraction of affective and prosodic information [14, 18-20]. These differences may reflect the lack of auditory spatial processing tasks among the paradigms used during construction of the MMP1 atlas.

### Orbitofrontal Cortex

Another example involves the orbitofrontal cortex. In the present synthesis, the parcel corresponding to posterior orbitofrontal cortex (pOFC) is associated primarily with sensory-specific motivational signals such as food-related reward [21], whereas the more anterior orbitofrontal cortex is more frequently associated with valuation of abstract stimuli, including monetary or aesthetic rewards [22]. Evidence from cortical stimulation studies reporting olfactory or somatosensory percepts following stimulation of posterior orbitofrontal sites [23], together with functional imaging studies examining valuation of food versus non-food items [24], suggests that the functional boundary between these regions may lie anterior to the border suggested by the MMP1 atlas.

### Gustatory Cortex and POI2

The synthesis highlights cases in which a single MMP1 parcel appears to encompass more than one literature-defined cortical field. For example, the parcel labeled POI2 in the MMP1 atlas appears to include two functionally related but distinct territories: a dorsal region corresponding to a primary gustatory and viscerosensory cortex (G1) and a ventral region corresponding to a more associative gustatory-olfactory cortex (G2). This distinction is supported primarily by depth-electrode recordings and stimulation studies within the insula [25-27], modalities that are less readily captured by conventional fMRI paradigms.

### Face-Selective Cortex

Similarly, the parcel labeled fusiform face complex (FFC) in the MMP1 atlas encompasses two face-selective regions that are consistently distinguished in the literature: a posterior occipital face area (OFA), associated with recognition of individual facial features, and a more anterior fusiform face area (FFA), involved in processing spatial relationships among facial features [28, 29]. In BEAP, the boundary between these territories was demarcated using a probabilistic functional atlas derived from fMRI contrasts between faces and objects [30].

### Supplementary Motor Area

In the MMP1 atlas, the supplementary motor area (SMA) is primarily associated with the SCEF parcel (supplementary and cingulate eye fields), with possible extension into the neighboring SFL parcel (superior frontal language area). Applying the present curation methodology, however, these parcels were subdivided into anterior and posterior components corresponding to anterior SMA (aSMA) and posterior SMA (pSMA). Although both subregions are broadly implicated in sequential and bilateral premotor planning [31], the functional implementation of these processes appears to differ across the two territories. Consequently, lesions to the anterior and posterior divisions are associated with distinct clinical manifestations. The posterior SMA is more frequently associated with control of trunk and limb movements, whereas the anterior SMA is more commonly linked to control of facial, ocular, laughter, and speech-related motor patterns. This distinction parallels a similar division of labor within the lateral premotor cortex, where dorsal premotor cortex (PMDc) is preferentially involved in body movement planning and ventral premotor cortex (PMv) in planning movements of the face and mouth.

### Auditory MBelt and PAC-R

The literature synthesis motivates reconsideration of several parcel labels proposed in the MMP1 atlas. One example concerns the auditory parcel labeled MBelt, which the MMP1 atlas identifies as homologous to the medial belt region in monkeys based on high-field fMRI evidence [32]. In contrast, much of the broader human auditory literature associates the region located on anterior Heschl's gyrus with the rostral primary auditory cortex (core area R), corresponding to the PAC-R designation used in BEAP [11, 33-42].

### VIP and Human Specialization

Another example involves the parcel labeled VIP in the MMP1 atlas, intended to imply homology with monkey area VIP based on a surface-based registration study [43]. That study, however, did not explicitly address human-monkey homology. BEAP corroborates VIP-like motion sensitivity in this region, corresponding to the BEAP label dIPSm [44-46]. However, motion processing in dIPSm appears more complex in humans [46], suggesting possible human-specific specialization. Moreover, neighboring regions, including dIPSa (MMP1 parcels 7AL and 7PC) and QCC (MMP1 parcels IP1 and IP2), also exhibit motion sensitivity and additional VIP-related properties. Both monkey VIP and human dIPSa respond to visual stimuli near the body [44, 47], and both monkey VIP and human QCC are implicated in quantity estimation [48-52]. The BEAP synthesis therefore raises the possibility that these regions represent human-specific derivatives of a VIP-like precursor, making a one-to-one homology assignment problematic.

### Perisylvian Language Area (PSL)

Based on an fMRI contrast between story comprehension and mental calculation, the MMP1 atlas proposed a language-related parcel labeled the perisylvian language area (PSL), which was described as a previously unreported cortical territory. The present synthesis, however, finds substantial convergence between PSL and the sylvian-parietal temporal area (Spt), a region extensively described in prior work as active during both speech perception and speech production [53-55] and strongly associated with speech repetition [56].

#### References:

1. Glasser, M.F., et al., *A multi-modal parcellation of human cerebral cortex.* Nature, 2016. **536**(7615): p. 171-178.
2. Fischl, B., et al., *Cortical folding patterns and predicting cytoarchitecture.* Cerebral cortex, 2008. **18**(8): p. 1973-1980.
3. Geyer, S., *The microstructural border between the motor and the cognitive domain in the human cerebral cortex.* 2012.

4. Vorobiev, V., et al., *Parcellation of human mesial area 6: cytoarchitectonic evidence for three separate areas.* European Journal of Neuroscience, 1998. **10**(6): p. 2199-2203.
5. Van der Zwaag, W., et al., *Where sound position influences sound object representations: a 7-T fMRI study.* Neuroimage, 2011. **54**(3): p. 1803-1811.
6. Smith, K.R., et al., *Auditory spatial and object processing in the human planum temporale: no evidence for selectivity.* Journal of cognitive neuroscience, 2010. **22**(4): p. 632-639.
7. Warren, J.D., et al., *Perception of sound-source motion by the human brain.* Neuron, 2002. **34**(1): p. 139-148.
8. Derey, K., et al., *Opponent coding of sound location (azimuth) in planum temporale is robust to sound-level variations.* Cerebral Cortex, 2016. **26**(1): p. 450-464.
9. Brunetti, M., et al., *Human brain activation during passive listening to sounds from different locations: an fMRI and MEG study.* Human brain mapping, 2005. **26**(4): p. 251-261.
10. von Kriegstein, K., et al., *Neural representation of auditory size in the human voice and in sounds from other resonant sources.* Current Biology, 2007. **17**(13): p. 1123-1128.
11. Moerel, M., F. De Martino, and E. Formisano, *Processing of natural sounds in human auditory cortex: tonotopy, spectral tuning, and relation to voice sensitivity.* Journal of Neuroscience, 2012. **32**(41): p. 14205-14216.
12. Formisano, E., et al., *" Who" is saying" what"? Brain-based decoding of human voice and speech.* Science, 2008. **322**(5903): p. 970-973.
13. Roswandowitz, C., et al., *Obligatory and facultative brain regions for voice-identity recognition.* Brain, 2018. **141**(1): p. 234-247.
14. Perrodin, C., et al., *Who is that? Brain networks and mechanisms for identifying individuals.* Trends in cognitive sciences, 2015. **19**(12): p. 783-796.
15. Sjerps, M.J., et al., *Speaker-normalized sound representations in the human auditory cortex.* Nature communications, 2019. **10**(1): p. 2465.
16. Lachaux, J.-P., et al., *A blueprint for real-time functional mapping via human intracranial recordings.* PLoS One, 2007. **2**(10): p. e1094.
17. Ahrens, M.-M., et al., *Gender differences in the temporal voice areas.* Frontiers in neuroscience, 2014. **8**: p. 228.
18. Sander, D., et al., *Emotion and attention interactions in social cognition: brain regions involved in processing anger prosody.* Neuroimage, 2005. **28**(4): p. 848-858.
19. Özdemir, E., A. Norton, and G. Schlaug, *Shared and distinct neural correlates of singing and speaking.* Neuroimage, 2006. **33**(2): p. 628-635.
20. Tang, C., L. Hamilton, and E. Chang, *Intonational speech prosody encoding in the human auditory cortex.* Science, 2017. **357**(6353): p. 797-801.
21. Rolls, E., *Functions of the orbitofrontal and pregenual cingulate cortex in taste, olfaction, appetite and emotion.* Acta Physiologica Hungarica, 2008. **95**(2): p. 131-164.
22. Rolls, E.T. and F. Grabenhorst, *The orbitofrontal cortex and beyond: from affect to decision-making.* Progress in neurobiology, 2008. **86**(3): p. 216-244.

23. Fox, K.C., et al., *Changes in subjective experience elicited by direct stimulation of the human orbitofrontal cortex.* Neurology, 2018. **91**(16): p. e1519-e1527.
24. McNamee, D., A. Rangel, and J.P. O'doherty, *Category-dependent and category-independent goal-value codes in human ventromedial prefrontal cortex.* Nature neuroscience, 2013. **16**(4): p. 479-485.
25. Isnard, J., et al., *Clinical manifestations of insular lobe seizures: a stereo-electroencephalographic study.* Epilepsia, 2004. **45**(9): p. 1079-1090.
26. Li, Y., et al., *Responses of chemosensory perception to stimulation of the human brain.* Annals of Neurology, 2023. **93**(1): p. 175-183.
27. Ostrowsky, K., et al., *Functional mapping of the insular cortex: clinical implication in temporal lobe epilepsy.* Epilepsia, 2000. **41**(6): p. 681-686.
28. Nakamura, K. and K. Kubota, *The primate temporal pole: its putative role in object recognition and memory.* Behavioural brain research, 1996. **77**(1-2): p. 53-77.
29. Pinsk, M.A., et al., *Neural representations of faces and body parts in macaque and human cortex: a comparative FMRI study.* Journal of neurophysiology, 2009. **101**(5): p. 2581-2600.
30. Zhen, Z., et al., *Quantifying interindividual variability and asymmetry of face-selective regions: a probabilistic functional atlas.* Neuroimage, 2015. **113**: p. 13-25.
31. Nachev, P., C. Kennard, and M. Husain, *Functional role of the supplementary and pre-supplementary motor areas.* Nat Rev Neurosci, 2008. **9**(11): p. 856-69.
32. Moerel, M., et al., *Using high spatial resolution fMRI to understand representation in the auditory network.* Progress in neurobiology, 2020. **207**: p. 101887.
33. Saenz, M. and D.R. Langers, *Tonotopic mapping of human auditory cortex.* Hearing research, 2014. **307**: p. 42-52.
34. Langers, D.R., *Assessment of tonotopically organised subdivisions in human auditory cortex using volumetric and surface-based cortical alignments.* Human brain mapping, 2014. **35**(4): p. 1544-1561.
35. Da Costa, S., et al., *Tonotopic gradients in human primary auditory cortex: concurring evidence from high-resolution 7 T and 3 T fMRI.* Brain Topogr, 2015. **28**(1): p. 66-9.
36. van Dijk, P. and D.R. Langers, *Mapping tonotopy in human auditory cortex*, in *Basic aspects of hearing: Physiology and perception*. 2013, Springer. p. 419-425.
37. Striem-Amit, E., U. Hertz, and A. Amedi, *Extensive cochleotopic mapping of human auditory cortical fields obtained with phase-encoding FMRI.* PloS one, 2011. **6**(3): p. e17832.
38. Dick, F., et al., *In vivo functional and myeloarchitectonic mapping of human primary auditory areas.* Journal of Neuroscience, 2012. **32**(46): p. 16095-16105.
39. Formisano, E., et al., *Mirror-symmetric tonotopic maps in human primary auditory cortex.* Neuron, 2003. **40**(4): p. 859-869.
40. Talavage, T.M., et al., *Tonotopic organization in human auditory cortex revealed by progressions of frequency sensitivity.* Journal of neurophysiology, 2004. **91**(3): p. 1282-1296.
41. Humphries, C., E. Liebenthal, and J.R. Binder, *Tonotopic organization of human auditory cortex.* Neuroimage, 2010. **50**(3): p. 1202-1211.

42. Petkov, C.I., et al., *Functional imaging reveals numerous fields in the monkey auditory cortex.* PLoS biology, 2006. **4**(7): p. e215.
43. Van Essen, D.C., et al., *Cortical parcellations of the macaque monkey analyzed on surface-based atlases.* Cerebral cortex, 2012. **22**(10): p. 2227-2240.
44. Bremmer, F., et al., *The representation of movement in near extra-personal space in the macaque ventral intraparietal area (VIP).* EXPERIMENTAL BRAIN RESEARCH SERIES, 1997. **25**: p. 619-630.
45. Colby, C.L., J.-R. Duhamel, and M.E. Goldberg, *Ventral intraparietal area of the macaque: anatomic location and visual response properties.* Journal of neurophysiology, 1993. **69**(3): p. 902-914.
46. Orban, G.A., et al., *Mapping the parietal cortex of human and non-human primates.* Neuropsychologia, 2006. **44**(13): p. 2647-2667.
47. Huang, R.-S., et al., *Mapping multisensory parietal face and body areas in humans.* Proceedings of the National Academy of Sciences, 2012. **109**(44): p. 18114-18119.
48. Wang, L., et al., *Representation of numerical and sequential patterns in macaque and human brains.* Current Biology, 2015. **25**(15): p. 1966-1974.
49. Dastjerdi, M., et al., *Numerical processing in the human parietal cortex during experimental and natural conditions.* Nature communications, 2013. **4**(1): p. 2528.
50. Rusconi, E., M. Turatto, and C. Umilta, *Two orienting mechanisms in posterior parietal lobule: an rTMS study of the Simon and SNARC effects.* Cognitive Neuropsychology, 2007. **24**(4): p. 373-392.
51. Cappelletti, M., et al., *rTMS over the intraparietal sulcus disrupts numerosity processing.* Experimental Brain Research, 2007. **179**: p. 631-642.
52. Montojo, C.A. and S.M. Courtney, *Differential neural activation for updating rule versus stimulus information in working memory.* Neuron, 2008. **59**(1): p. 173-182.
53. Wise, R.J., et al., *Separate neural subsystems withinWernicke's area'.* Brain, 2001. **124**(1): p. 83-95.
54. Hickok, G., et al., *Auditory–Motor Interaction Revealed by fMRI: Speech, Music, and Working Memory in Area Spt.* Journal of Cognitive Neuroscience, 2003. **15**(5): p. 673-682.
55. Hickok, G. and D. Poeppel, *The cortical organization of speech processing.* Nature reviews neuroscience, 2007. **8**(5): p. 393-402.
56. Buchsbaum, B.R., et al., *Conduction aphasia, sensory-motor integration, and phonological short-term memory–an aggregate analysis of lesion and fMRI data.* Brain and language, 2011. **119**(3): p. 119-128.
