## Supplementary material for "The Brain Encyclopedia Atlas Project (BEAP): A Literature-Synthesis-Derived Functional Atlas of the Human Brain": 5 supplemental sections: SupplementaryMaterial5.pdf

**Table 3. Reproducibility of figure-to-parcel assignments across cortical systems**

| Representative BEAP cortices | Studies re-curated (n) | Exact parcel agreement (%) | Adjacent parcel disagreement (%) | Different cortical system (%) |
|---|---|---|---|---|
| aIns | 5 | 5 | 0 | |
| OPA | 5 | 4 | 1 | |
| sBroca | 5 | 5 | 0 | |
| PP | 5 | 5 | 0 | |
| FPm | 5 | 4 | 1 | |
| aITL | 5 | 5 | 0 | |
| EFA | 5 | 5 | 0 | |
| QCC | 5 | 5 | 0 | |
| RSC | 5 | 4 | 1 | |
| pMCC | 5 | 5 | 0 | |
| OFCl | 5 | 4 | 1 | |
| MTc | 5 | 5 | 0 | |
| Spt | 5 | 5 | 0 | |
| dIPSm | 5 | 4 | 1 | |
| TPJp | 5 | 4 | 1 | |
| PMDr | 5 | 5 | 0 | |
| S1-3b | 5 | 5 | 0 | |
| pIns | 5 | 5 | 0 | |
| aSMA | 5 | 5 | 0 | |
| V3 | 5 | 5 | 0 | |
| Total | **100** | **94** | 6 | **0** |

To evaluate procedural stability, a subset of 100 studies (5 studies of 20 randomly selected areas) was independently re-curated approximately one year after the initial annotation while blinded to prior assignments (supplementary material 5). Of the 100 re-evaluated studies, only in 6 cases the activation was shown in an adjacent parcel upon second inspection, and in no study was the activation re-attributed to a non-neighboring parcel.
